# Comparative Analysis Reveals Host Species-Dependent Diversity Among 16 Virulent Bacteriophages Isolated Against Soybean *Bradyrhizobium* spp

**DOI:** 10.1101/2025.10.06.680108

**Authors:** Emily A. Morgese, Barbra D. Ferrell, Spencer C. Toth, Shawn W. Polson, K. Eric Wommack, Jeffry J. Fuhrmann

**Affiliations:** Department of Plant and Soil Sciences, University of Delaware, Newark, DE 19716, USA; Department of Biological Sciences, University of Delaware, Newark, DE 19713, USA; Department of Computer and Information Sciences, University of Delaware, Newark, DE 19713, USA; Microbiology Graduate Program, University of Delaware, Newark, DE 19713, USA

**Keywords:** *Bradyrhizobium*, soybean, bacteriophages

## Abstract

Phages play a role in shaping ecosystems by controlling host abundance via cell lysis, driving host evolution via horizontal gene transfer, and promoting nutrient cycling. The genus *Bradyrhizobium* includes bacteria able to symbiotically nodulate the roots of soybean (*Glycine max*), providing the plant with a direct source of biologically fixed nitrogen. Optimizing this symbiosis can minimize the use of nitrogen fertilizers and make soybean production more sustainable. Phages targeting *Bradyrhizobium* may modify their hosts’ genotype, alter phenotypic traits such as symbiotic effectiveness, and mediate competition among strains for nodulation sites. Sixteen phages were isolated against *B. elkanii* strains USDA94 and USDA31, and *B. diazoefficiens* strain USDA110. Comparative analyses revealed host species-dependent diversity in morphology, host range, and genome composition, leading to the identification of three previously undescribed phage species. Remarkably, all *B. elkanii* phages shared a siphophage morphology and formed a single species with >97% nucleotide identity, even when isolated from farms separated by up to ∼70 km, suggesting genomic stability across geographic scales. In contrast, phages isolated against *B. diazoefficiens* displayed podophage-like morphology, greater genetic diversity, and divided into two distinct species. Although no phages were recovered against *B. japonicum* strains or native Delaware *Bradyrhizobium* isolates tested, some Delaware isolates showed susceptibility during the host range assay. The phage genomes demonstrated features predicting phenotypes. Terminase genes predicted headful packaging among the phages which is critical for generalized transduction. The *B. elkanii* phages all carried tmRNA genes capable of recruiting stalled ribosomes and both phage groups carried DNA polymerase A indicating greater control of phage genome replication. State-of-the-art structural annotation revealed a tail fiber gene within a phage genome having the highest proportion (80.77%) of unknown genes. Together this work expands the limited knowledge available on soybean *Bradyrhizobium* phage ecology and genomics.

## 1. Introduction

Soybean (*Glycine max*) is a cornerstone of global agriculture, with an estimated 420.9 million metric tons of seed produced worldwide in 2024 [1]. According to the same report, the United States accounted for over 25% of that production, ranking it second globally. Soy is valued for its high protein content, the highest among all plant-based food crops, and its wide range of applications, including animal feed, human nutrition, biofuel production, and industrial uses [2,3]. The high nutritional value and affordability of soy make it essential in addressing global food security.

Producing protein-rich seed places a high nitrogen (N) requirement on the soybean plant. A portion of this N is provided through symbiotic root-nodulating *Bradyrhizobium* bacteria[4]. However, this requirement is increasingly supplemented by applying synthetic N fertilizers to the soil [5], the production of which relies on fossil fuels and contributes significantly to CO_2_ emissions to the atmosphere [6]. Farmers often apply these fertilizers in amounts exceeding the soil N use efficiency of the plant; the unassimilated N, largely in the form of nitrate (NO_3_^−^), is prone to leaching or denitrification.[7]. Elevated nitrate levels in ground and surface waters can have adverse human health effects when consumed and often lead to algal blooms that disrupt aquatic ecosystems [8,9]. Additionally, nitrous oxide (N_2_O) produced via denitrification contributes to global warming and stratospheric ozone depletion [10]. Inoculating soybean crops with highly efficient nitrogen-fixing bradyrhizobia strains can improve yield without supplemental nitrogen fertilizers [11]. Bradyrhizobia reduce atmospheric nitrogen (N2) to ammonia (NH3) within root nodules where it is directly assimilated by the plant, thereby reducing the likelihood of nitrate pollution and greenhouse gas emissions [12]. Enhancing biological N fixation by *Bradyrhizobium* can lower production costs by decreasing synthetic fertilizer application and improve the sustainability of soybean cultivation by reducing environmental impacts.

Bacteriophages (phages), viruses that infect bacteria, are the most abundant biological entities on Earth. Current estimates suggest that there are upwards of 10^31^ phages in the biosphere, outnumbering their hosts by an order of magnitude [13,14]. They play a critical role in microbial ecosystems by controlling bacterial populations through lysis, facilitating horizontal gene transfer, and contributing to nutrient cycling [15,16]. Although numerous studies have examined various aspects of the soybean– *Bradyrhizobium* symbiosis, relatively few have examined phages that infect soybean bradyrhizobia. Most of these studies have isolated and characterized a limited number of virulent *Bradyrhizobium* phages, focusing primarily on phenotypic traits such as morphology, host range, and infection [17–24]. Notably, one study found that a virulent phage altered nodulation competition between two bradyrhizobia strains under gnotobiotic conditions, significantly reducing nodulation by a susceptible strain [20]. This phenomenon of phages negatively impacting nodulation was also reported for *B. japonicum* lysogens [23]. The negative impact of phages on nodulation could result from spontaneous prophage production from lysogenic strains [43]. Recent studies found that a majority of soybean bradyrhizobial isolates spontaneously produce temperate bacteriophages under laboratory conditions [25]. Time course experiments showed that phages were spontaneously produced throughout host growth, and genome sequencing indicated spontaneously produced phages may be capable of gene transduction [24]. However, few comprehensive genomic comparisons of *Bradyrhizobium* phages, temperate or virulent, have been reported.

Phage genomes are highly diverse and often mosaic, composed largely of genes of unknown function, presenting challenges for functional annotation and comparative genomic analyses [26]. Phage taxonomy has evolved from morphology-based classification to genome-based criteria, with current standards designating phages with ≥95% average nucleotide identity (ANI) as belonging to the same species, and those sharing ≥99.5% belonging to the same subspecies [27–30], or genomovar [30]. Morphological traits, such as tail structure and capsid volume, are often correlated with genome size and host specificity, making them useful indicators of phage behavior and ecological interactions [31–33]. Understanding phage genomic and phenotypic features can enable predictions of life history traits and host interactions in environments such as soils and the plant rhizosphere.

Despite ecological and agricultural significance of soybean *Bradyrhizobium* spp., little is known about the genomic diversity of associated virulent phages. This study addressed this knowledge gap through sequencing the genomes of sixteen novel virulent phages infecting *B. elkanii* and *B. diazoefficiens* hosts isolated from Delaware soybean fields. These phages were phenotypically assessed for morphology and host range, and their genomes were subjected to comparative analysis. The possible role of geography on phage diversity was also examined. Together, these findings significantly expand current knowledge of virulent phages infecting soybean bradyrhizobia and lay the foundation for future investigations into their ecological roles in the *Bradyrhizobium*–soybean symbiosis and their potential applications in promoting sustainable soybean production.

## 2. Materials and Methods

### 2.1. Lytic Phage Isolation

Soil samples were collected from 10 soybean fields across Kent and Sussex counties in Delaware, USA, and enriched for lytic phage activity against 10 *Bradyrhizobium* strains from the University of Delaware *Bradyrhizobium* Culture Collection (UDBCC; [25,34]), i.e., 100 soil-host combinations in total (Figure 1). The strains included five USDA reference strains from the USDA-ARS National Rhizobium Germplasm Collection (Beltsville, MD; https://www.ars-grin.gov/Rhizobium) and five strains isolated from Delaware soils. To isolate phage, 25 g of a given soil was added to 100 mL of sterile SM buffer (pH 8.0; 50 mM Tris HCl, 100 mM NaCl, 20 mM MgSO_4;_ Fisher Scientific, Waltham, MA), shaken (150 rpm) for one hour to release phages into suspension, and then centrifuged (1000 RCF, 10 min) to pellet soil particles. The supernatant was collected and filtered using 0.2 μm Whatman Anotop® filters (GE Healthcare UK Limited, Buckinghamshire, United Kingdom) to remove cells. Five milliliters of filtered soil extract was added to 20 mL of a logarithmically growing *Bradyrhizobium* culture in Modified Arabinose Gluconate (MAG) medium (ATCC medium 2233) and incubated with shaking (150 rpm) at ambient temperature (∼24 °C) for 72-120 hours to allow for phage replication. Following incubation, cultures were centrifuged and filtered as previously described. A 10 μL aliquot of the filtrate was spotted onto a Petri plate containing 25 mL of MAG base agar (1.5% agar; Becton Dickinson, Franklin Lakes, NJ) overlaid with 5 mL of MAG soft agar (0.7% agar) seeded with exponentially growing cells (approximately 10^7^ cells mL–^1^ soft agar) of the enrichment strain. Plates were incubated at 30 °C for 72-120 hours to allow clearings to become apparent. Phage isolation and purification were performed by quadrant-streaking a loopful of agar from clearings onto a 1.5% MAG agar base and overlaying with a 0.7% MAG soft agar layer containing the same host strain used originally. Plates were incubated as previously described, and well-separated plaques were used for further isolation and purification. This latter procedure was performed a total of three times to ensure the purity of the lytic phage isolates. Purified phage isolates were mixed 1:1 with 30% v/v glycerol (Fisher Scientific) and stored at −80°C.

**Figure 1.**
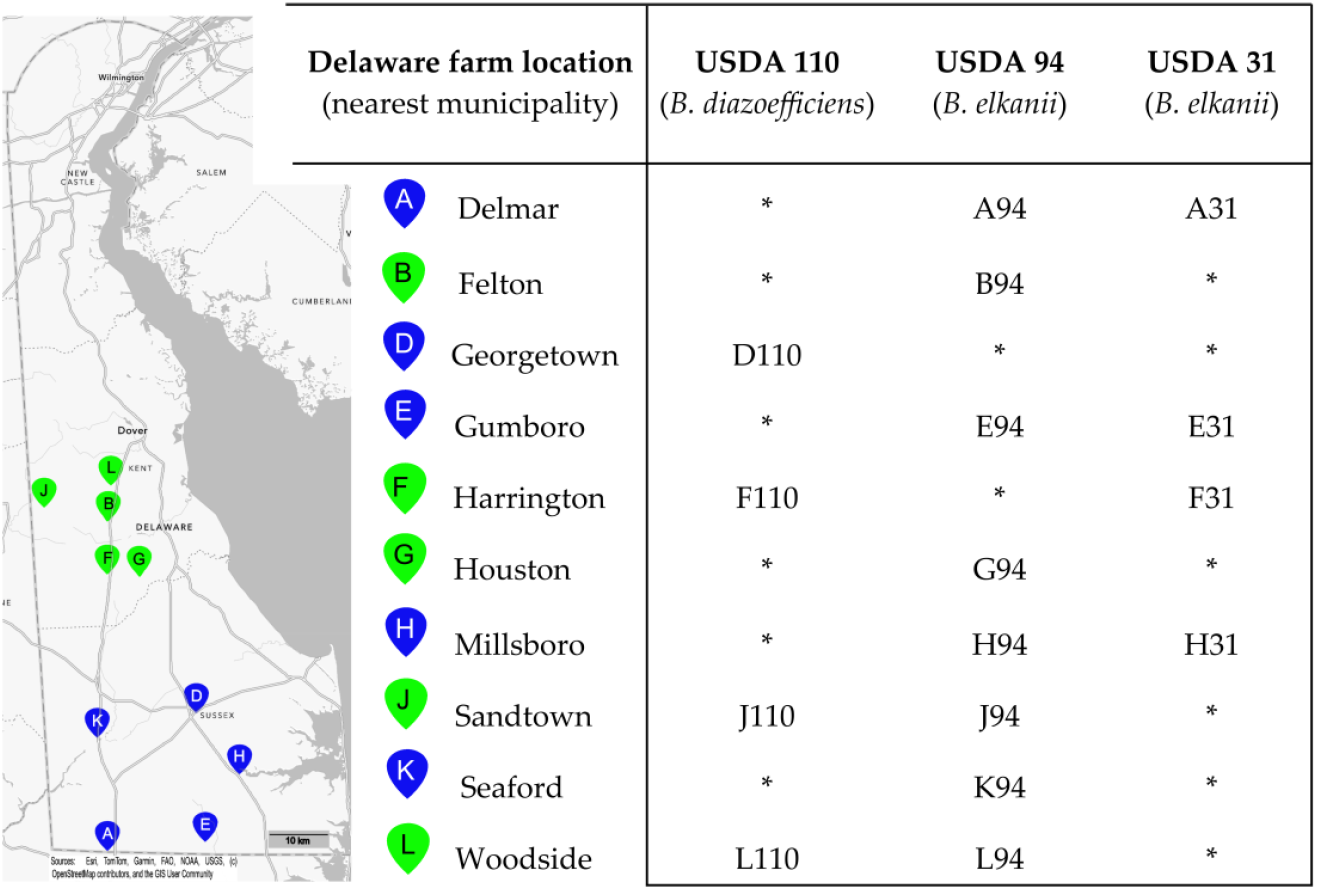
The map on the left portion of the figure shows the locations of 10 soybean fields in Delaware, USA, from which soils were sampled for virulent phage isolation. The pin letter codes correspond to the sample site, and the colors indicate the county in which the site was located (green: Kent County; blue: Sussex County). Matrix (at right) defines the *Bradyrhizobium* isolation host strain and soil sampling location for each of the 16 virulent bacteriophages, with phage names derived from the letter code of the sampling site followed by a numerical code indicating the *Bradyrhizobium* strain against which they were isolated. No phages were isolated against seven additional strains tested: *B. japonicum* strains S06G, K03C, and S04E (all native Delaware strains) or USDA 123; *B. diazoefficiens* Delaware strains K09F and N03H; *B. elkanii* strain USDA 46. ^*^No phage was isolated from the soil-strain combination.

### 2.2 Lysate Preparation and Titering

Lysates were prepared in MAG broth culture by adding a 10-uL aliquot of phage stock to 50 mL of a logarithmically growing culture of the *Bradyrhizobium* strain initially used for enrichment and isolation. An exception was made for three phages initially isolated against strain USDA 94. USDA 94 was previously shown to spontaneously produce phages in laboratory culture [25]. We observed during the current study that infection with three of the lytic phages isolated against USDA 94 (J94, K94, L94) apparently increased spontaneous phage production by this host. To prevent these temperate phages from interfering with the preparation of virulent phages, lysates of these three phages were instead prepared using an alternate susceptible host, Delaware-isolated *B. elkanii* strain K03D, which does not spontaneously produce phages [25]. The infected cultures were incubated at 30 °C for 18 hours with shaking (150 rpm). Following this overnight incubation, the culture was split by transferring 25 mL of the culture to a sterile flask, and 25 mL of fresh MAG broth was added to both the new and original flasks, doubling the total volume of the culture. These cultures were incubated overnight as previously described, centrifuged for 15 min at 10,000 RCF to pellet cells, and the combined supernatants were filtered through a 0.2 μm Whatman Anotop® filter. Phages in the filtrate were concentrated using 100 kDa Amicon® Ultra Centrifugal Filters (Millipore, Lexington, MA) at 4,000 RCF for 10 min to obtain a final volume of approximately 5 mL from the original 100 mL of culture.

To determine phage numbers in the lysates, 250 μL of lysate was collected onto a 0.02 μm Whatman® Anodisc membrane filter and stained with 200 μL of 2× SYBR Gold DNA stain (Invitrogen, Fisher Scientific) for 10 min. The filter was then washed twice with 500 μL of MSM buffer (pH 7.5; 1.2 g Tris-HCL, 4.68 g NaCl, 0.986 g MgSO_4_; Fisher Scientific) and transferred to a microscope slide spotted with 20 μL of antifade solution (0.1% p-phenylenediamine; Sigma-Aldrich, St. Louis, MO) to prevent photobleaching. An additional 20 μL of antifade solution was applied to the top of the filter followed by a coverslip. Virus-like particles (VLPs) were digitally imaged at 1000× magnification using an epifluorescence microscope equipped with a fluorescein (FITC) filter set. Image processing software (Serif PhotoPlus X8 v18.0.0.15) was used to superimpose a 5-by-5 grid of squares of known area onto each micrograph. For each phage, approximately five images were analyzed. For each image, VLP counts were taken for 10 randomly selected squares. The mean VLP counts from the five images were then averaged to estimate the concentration of VLPs in the original lysate.

### 2.3 Electron Microscopy of Phages

Transmission electron microscopy was used to examine the morphological characteristics of isolated phages. Concentrated phage lysates were imaged at the University of Delaware Bio-Imaging Center (Newark, DE) as previously described [25]. Capsid, collar, tail, and baseplate dimensions were measured using ImageJ Fiji 2.0.0 (National Institute of Health, Bethesda, MD) [35]. Capsid volumes were calculated using a formula for the volume of an ellipsoid: V = 4/3πa^2^c where V = volume of an ellipsoid, a = semi-axis of the capsid width (capsid width/2), and c = semi-axis of the capsid length (capsid length/2) [36]. Welch’s t-test was used to assess statistical differences in the mean measurements for each morphologic characteristic. Morphological measurements were also visualized as jittered boxplots using the ggplot2 package in RStudio [37].

### 2.4 Host Range Determination

The host range of 16 isolated phages was assessed using spot assay on overlay plates seeded with a subset of 12 *Bradyrhizobium* strains from the UDBCC representing four species that nodulate soybean (*B. diazoefficiens, B. elkanii, B. japonicum*, and *B. ottawaense*). Host cultures were grown in MAG broth at 30 °C with shaking (150 rpm) until they reached an optical density (OD_600_) of approximately 1.0. Cultures were then diluted back to an OD of 0.05 with fresh MAG and incubated for an additional 24 h until reaching mid-log phase (OD 0.4–0.6). On the day of plating, MAG containing 0.6% low-melt agar (IBI Scientific, Dubuque, IA) was melted and maintained at 40 °C in a water bath. For each overlay, 500 µL of mid-log host culture was mixed with 4.5 mL of molten soft agar and poured over solidified MAG base agar (1.5% agar; Becton Dickinson). These plates were allowed to solidify at room temperature for approximately 20 min. Phage suspensions were prepared by adding approximately 10 μL of phage stock to 1 mL of sterile MSM buffer. After the overlays had set, 10 µL of each phage suspension was spotted onto the surface of duplicate plates. Plates were left in a sterile laminar flow hood for approximately 1 h to allow the phages to absorb into the agar. The plates were then incubated at 30 °C for five days or until clear zones of lysis were visible.

### 2.5 DNA Extraction and Sequencing

Phage lysates having a concentration of at least 10^7^ VLPs mL–^1^ were used for DNA extraction. To remove bacterial DNA prior to phage DNA extraction, an 8*-*µL aliquot of DNase I solution (2.5 units/µL; Fisher Scientific) was added to 1 mL of concentrated phage lysate to obtain a final concentration of 20 units mL–^1.^ This mixture was incubated at room temperature for 15 min. The enzyme was then inactivated by incubating for 5 min at 75°C. A 16S rRNA PCR was performed on the DNase-treated lysates to check for bacterial contamination using forward and reverse primers (Integrated DNA Technologies (IDT), Coralville, IA) chosen from those described by Klindworth et al. [38], specifically S-D-Bact-0341-b-S-17 (5’-CCTACGGGNGGCWGCAG-3’) and S-D-Bact-0785-a-A-21 (5’-GACTACHVGGGTATCTAATCC-3’). The PCR amplicons were visualized via standard 1.3% agarose gel electrophoresis on a gel stained with 0.01% v/v Invitrogen SYBR Safe DNA gel stain (Fisher Scientific). Hi-Lo DNA Markers (Bionexus®, Oakland, CA) were used as the DNA ladder, and the gel was imaged using a SYBR Gold gel imaging filter. If the gel showed no evidence of bacterial DNA contamination (no bands matching the positive control; expected amplicon length = 462 bp), the sample was used for viral DNA extraction.

DNA extraction was performed using the Norgen Biotek® Phage DNA Isolation Kit (Norgen Biotek Corp., Thorold, Ontario) following the recommended manufacturer protocol with minor modifications. Specifically, a proteinase K step was added prior to the addition of isopropanol to promote the release of DNA from the viral capsids. A 4-µL aliquot of MP Biomedicals® Proteinase K (20 mg/mL) (MP Biomedicals, Santa Anna, CA) was added to the mixture and incubated for 30 min in a 55 °C water bath. The mixture was then incubated for an additional 15 min at 65 °C. Subsequently, the remainder of the manufacturer’s recommended protocol was followed. The released DNA was stored short-term (∼ 1 week) at −20 °C and long-term at −80 °C. DNA concentrations were measured using the Qubit® dsDNA High Sensitivity Quantification Assay (Fisher Scientific).

Illumina whole genome sequencing was performed by SeqCenter (Pittsburgh, PA). Sequencing libraries were prepared using the Illumina DNA Prep kit (Illumina, San Diego, CA) and custom IDT 10 bp unique dual indices (UDI). Sequencing was performed on an Illumina NextSeq 2000 sequencer, producing 2×151 bp paired end reads. Demultiplexing, quality control and adapter trimming was performed with bcl-convert (v3.9.3).

### 2.6 Assembly and Annotation of Lytic Phage Genomes

Illumina reads were assembled into contiguous sequences (contigs) using the SPAdes Genome Assembler (version 4.0.0) [25]. The contigs from each assembly that had the highest coverage and anticipated genome length (approximately 50 - 60 kb) were chosen as the viral contig. The assembled contigs were mapped back to the genome of their respective *Bradyrhizobium* isolation host using the Geneious Prime® mapping tool (version 2024.0.7) with default settings to screen for contaminating bacterial DNA and assess the potential presence of temperate phage sequences. This latter step was necessary because some soybean-associated *Bradyrhizobium* strains have been shown to spontaneously produce phages under laboratory conditions [25], introducing the possibility that recovered phage sequences may originate from induced temperate prophages rather than the virulent isolates. If a contig aligned to the host genome, its location was cross-referenced with predicted prophage regions [25] to determine if it corresponded to a putative prophage. Only those contigs that did not map back to the host genome were considered to represent virulent phage isolates.

Gene prediction and annotation were performed using the Pharokka Galaxy interface with default parameters (version 1.3.2 + Galaxy 0) [39], which employs Prodigal [40] as the gene caller and uses the Pharokka database (version 1.2.0) to assign annotations [41]. To facilitate comparative analysis, all the genomes were rearranged to start with the annotated large subunit terminase gene (*TerL*) and reoriented to have the same directionality [41]. Further annotation of the *TerL* gene was performed by identifying associated protein family (Pfam) domains [42], which can be used to predict the phage DNA packaging mechanism [43–45]. Additionally, further investigation of the *polA* genes was performed, specifically identifying the residue at the 762 (E. coli numbering) position in these genes as previously described [46]. Notably, the automatically assembled E94 genome was approximately 2.3 kb shorter than the other *B. elkanii* phages due to an apparent truncation of the annotated bifunctional primase-polymerase (prim/pol) gene. However, mapping the E94 sequencing reads back to the prim/pol genes from the other *B. elkanii* phages showed they spanned the entire length of the gene. To correct this assembly issue, the fragmented E94 prim/pol gene was manually reconstructed using the consensus sequence of the prim/pol genes found in the other 11 *B. elkanii* phage genomes. Hypothetical genes of interest were further investigated using structural homology searches to predict their potential function. Protein sequence structure was predicted in the AlphaFold server [47] and iterative structural homology searches with Foldseek [48] were performed against the AlphaFold/UniProt50 v4, BFVD 2023_02, and AlphaFold/Swiss-Prot v4 databases. Hits that had TM-scores above 0.5 were evaluated for functional annotation.

### 2.7 Taxonomic Classification

Phage taxa were first assessed using taxMyPhage v0.3.6 [49] which classifies dsDNA bacteriophages at the genus and species levels based on nucleotide similarity to ICTV-curated reference genomes. The tool was run with default settings. Genomes lacking significant matches were manually classified to higher taxonomic ranks following ICTV guidelines [50], based on genome composition (dsDNA) and tail morphology inferred from transmission electron microscopy.

### 2.8 Average Nucleotide Identity and Coverage

Pairwise average nucleotide identity (ANI) and corresponding genome coverage values were calculated for the 16 genomes using FastANI v1.33 [51]. For each pairing, ANI values, the number of aligned fragments, and the total number of query fragments were recorded. Genome coverage was calculated as: coverage = (aligned fragments/total fragments) × 100. Because FastANI only reports coverage in one direction (query vs. reference), reciprocal comparisons were averaged to compute a symmetric coverage matrix. A combined heatmap of ANI and coverage values were generated using Python v3.12 with the following libraries: pandas v2.2.1, NumPy v1.26.4, seaborn v0.13.2, and matplotlib v3.8.4.

### 2.9 Genome Maps and Alignments

Based on the ANI-determined grouping of the 16 phages into three species, the longest genome from each phage species was selected as the representative genome for visualization. Annotated linear genome maps depicting gene order, orientation, and predicted function were generated using Proksee [52]. Whole-genome alignments of phage sequences were visualized using a custom Python script that computes pairwise nucleotide identity in 50 bp windows across a multiple sequence alignment. An alignment was created for each ANI-determined phage group containing more than one phage, and the longest genome (representative genome) was selected as the reference sequence. The script calculates the proportion of matches between each query sequence and the reference. Nucleotide identity was plotted as Integrative Genomics Viewers (IGV)-style horizontal tracks, and genome-wide consensus identity was included to highlight conserved and divergent regions across the alignment, as well as gaps. To supplement the whole-genome alignments **C**linker (v0.0.31) alignments [53] and a pangenome analysis using Anvi’o (v7.1) [54,55] were used to observe the overall relationships among the 16 phages.

### 2.10 Geographical Analysis

A geographic analysis was performed to detect possible relationships between phage isolation location and ANI. Specifically, the Mantel test [56] for correlation between two distance matrices was performed using the SciKit-Bio stats package [57]. ANI distance was calculated as: (100 − ANI). The geographic distance was calculated using GeoPy by determining the distance in kilometers between two isolation site locations based on their latitude and longitude values [58]. The calculated ANI and geographic distances were used to populate two symmetric matrices which served as the input for the Mantel test. The Mantel test was performed on three subsets of data: all 16 phages, only those phages isolated against *B. elkanii*, and only those phages isolated against *B. diazoefficiens*.

## 3. Results

### 3.1 Lytic Phage Isolation

Ten soil samples from soybean farms in Delaware were tested against 10 *Bradyrhizobium* host strains for the presence of lytic phages. From the 100 pairings, 16 lytic phages were isolated against three host strains: *B. elkanii* strains USDA 31 and USDA 94, and *B. diazoefficiens* strain USDA 110 (Figure 1). All 10 soils yielded at least one phage isolate. Twelve phages infected a *B. elkanii* host (USDA 94 or 31), and four infected USDA110. Strain USDA 94 yielded twice the number of phages isolates as USDA 31. Only three farms yielded phages infecting both host species. No phages were isolated against any of the four *B. japonicum* hosts nor any of the native Delaware *Bradyrhizobium* strains tested.

### 3.2 Transmission Electron Microscopy and Phage Morphology

TEM imaging of the 16 phage isolates revealed distinct morphological differences between those isolated against a *B. elkanii* host versus those isolated against the *B. diazoefficiens* host. The 12 phages isolated against a *B. elkanii* host had a siphophage-like morphology with prolate capsids and long, non-contractile tails. In contrast, the four phages isolated against the *B. diazoefficiens* strain USDA 110 host had icosahedral heads and an apparently podophage-like morphology, although the presence of tail structures for J110 could not be definitively confirmed (Figure 2).

**Figure 2.**
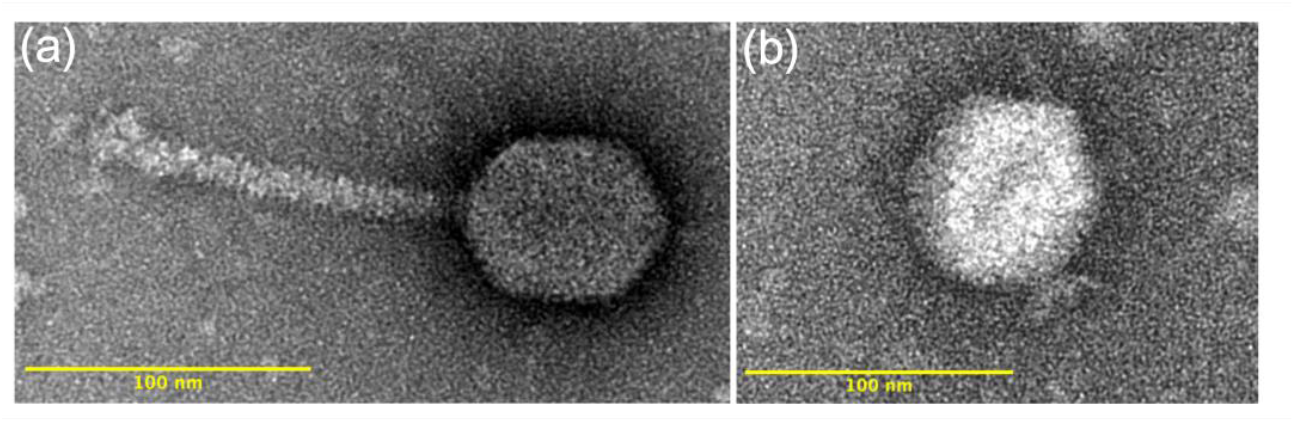
Representative transmission electron images of virulent phages isolated from Delaware soils against (a) *Bradyrhizobium elkanii* strains USDA 31 and USDA 94 (phage A31 shown) and (b) *B. diazoefficiens* strain USDA 110 (phage D110 shown).

Capsid dimensions and volumes were compared across phage isolates within both host species and host strain (Table S1, Figure S1). Mean lengths and diameters of the prolate capsids of *B. elkanii* phages did not differ significantly (p > 0.05), with average lengths of 71.5 nm and 72.2 nm and diameters of 58.1 nm and 59.3 nm for USDA 31 and USDA 94, respectively. The mean capsid diameter of *B. diazoefficiens* (USDA 110) phages (67.3 nm) was significantly greater (p = 1.18E–27) than the *B. elkanii* phages (58.9 nm). Additionally, the mean capsid length of the *B. elkanii* phages (72.0 nm) was significantly greater (p = 1.86E–10) than the *B. diazoefficiens* phages (67.9 nm). The mean capsid volumes of USDA 31 and USDA 94 (*B. elkanii*) phages were not significantly different; however, there was a significant difference (p = 4.18E–15) in mean capsid volumes between the *B. elkanii* phages (1.32E5 nm^3^) and the *B. diazoefficiens* phages (1.63E5 nm^3^). Mean tail lengths and diameters for the *B. elkanii* phages were 127.1 nm and 12.5 nm, respectively, and did not differ significantly (p > 0.05) between the two host strains. The podo-like tails of the phages isolated against *B. diazoefficiens* strain USDA 110 were often positionally or otherwise obscured and therefore difficult to measure reliably; however, they averaged approximately 17 nm in length.

### 3.3 Host Range

Of the 12 *Bradyrhizobium* strains tested for susceptibility to lytic phages, seven were susceptible to at least one of the 16 phage isolates (Figure 3). Lysis outcomes were binned into one of three categories based on spot assay appearance: complete lysis (clear spot), partial lysis (turbid spot), or no lysis (no clearing) (Figure S2). *B. diazoefficiens* strains USDA 110 and S13E were susceptible to all four phages originally isolated against USDA 110; conversely USDA 122 from the same species was resistant. Two *B. elkanii* strains, USDA 130 and S05J, exhibited resistance to all 16 phages. Strain S07J (*B. elkanii*) displayed partial lysis by the eight phages isolated against USDA 94 (Figure S2) and was resistant to the remaining phages. Notably, strains S01C, USDA 31, USDA 76, and USDA 94 (all *B. elkanii)* were susceptible to all 12 phages initially isolated against that host species, exhibiting complete lysis, with the exception that USDA 94 showed only partial lysis with phage F31. Strains USDA 123 (*B. japonicum*) and S15A (*B. ottawaense*) were resistant to infection by all 16 phages.

**Figure 3.**
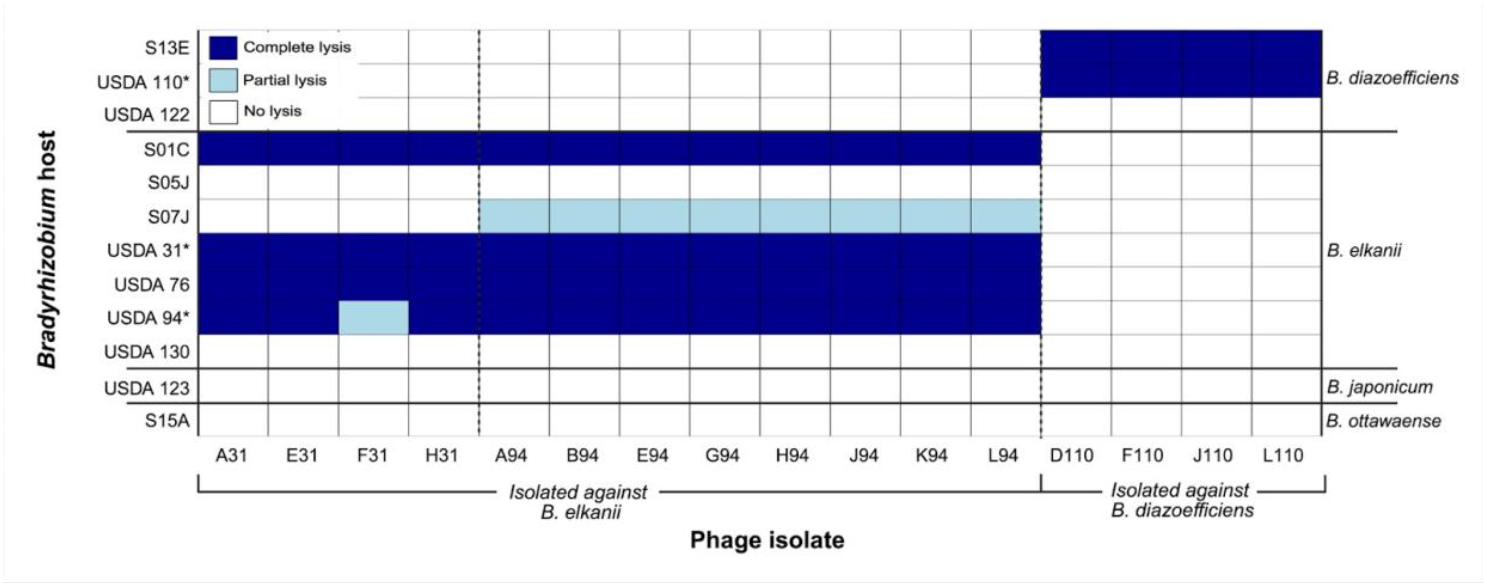
Host range of 16 virulent *Bradyrhizobium* phages isolated from Delaware soils (x-axis) against 12 soybean *Bradyrhizobium* strains representing four species (strains on left y-axis, corresponding species on right y-axis). The phage isolates are grouped according to the host isolation strain: *B. elkanii* strains USDA 31 and USDA 94, and *B. diazoefficiens* strain USDA 110. Lysis outcomes were classified into three categories based on spot assay appearance: complete lysis (clear zone), partial lysis (turbid zone), or no lysis (no clearing).

^*^ *Bradyrhizobium* strain used for the isolation of lytic phages.

### 3.4 Genome Sequencing

Genome length and %GC content tentatively separated the phage isolates into three groupings (Table S2), which were subsequently confirmed by ANI analysis. The *B. elkanii* phages were very similar in genome length and G+C content, whereas *B. diazoefficiens* phages formed two groups. The average genome length of *B. elkanii* phages was 62,920 bp, compared with 53,037 bp for the *B. diazoefficiens* phages; however, there was approximately a 15,000 bp difference between the size of the J110 genome (41,973 bp) and the mean genome length for the other three *B. diazoefficiens* phages (56,725 bp). The *B. elkanii* phages had a greater mean G+C content (67.0%) than the *B. diazoefficiens* phages (47.4%). Confirming the uniqueness of J110, its G+C content was 58.9% compared to a mean G+C content of 43.6% for the other three *B. diazoefficiens* phages.

### 3.5 Taxonomic Classification

Taxonomic analysis did not yield genus or species level matches for any of the phages, suggesting they represent novel lineages. Based on their tailed morphology as confirmed by TEM and shared double-stranded DNA genome structure, the phages were manually classified under the current ICTV taxonomy as Realm *Duplodnaviria*, Kingdom *Heunggongvirae*, Phylum *Uroviricota*, and Class *Caudoviricetes*.

### 3.5 Average Nucleotide Identity

All the *B. elkanii* phages shared an ANI above 95% at a coverage of at least 85%, with the highest values observed between E31 and E94 (100.0% ANI, 95.00% coverage) and H31 and H94 (100.0% ANI, 95.24% coverage) (Figure 4). The *B. elkanii* and *B. diazoefficiens* phages shared below 80% ANI with each other. The *B. diazoefficiens* phages were more diverse, with D110, F110, and L110 sharing between 95% and 97% identity, but with corresponding coverages between 84% and 94%. Conversely, J110 shared less than 80% ANI with the other 15 phages. Based on these ANI scores, all the phages isolated against *B. elkanii* grouped into a single phage species, whereas those isolated against *B. diazoefficiens* represented two species, with D110, F110, and L110 comprising one species and J110 being a separate unique species.

**Figure 4.**
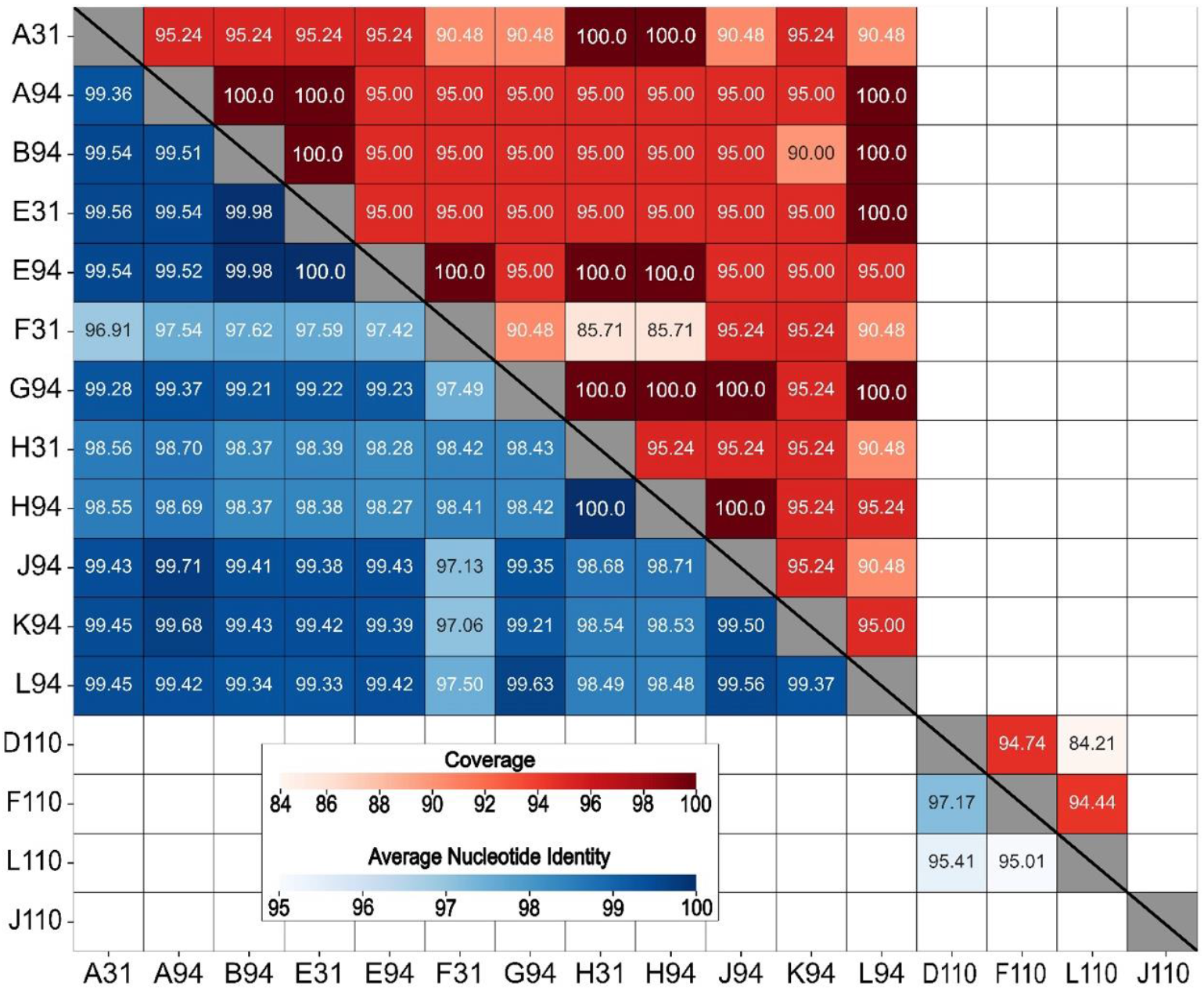
Heatmap of whole genome average nucleotide identity (ANI, blue-shaded lower half) and coverage scores (red-shaded upper half) for 16 virulent bacteriophages isolated against *Bradyrhizobium* phages. ANI and coverage are not displayed for pairings with ANI < 80% (unlabeled boxes).

### 3.6 Gene Prediction and Annotation

The total number of ORFs predicted per genome varied slightly across the 16 sequenced genomes (Table S2). The average number of genes predicted for the *B. elkanii* and *B. diazoefficiens* phages was 99.9 and 96.5, respectively. However, the latter value increased to 102.7 genes when J110 outlier (78 genes) was omitted. Overall, approximately 24% of the genes predicted for each genome were annotated as a protein of known function, with the remainder denoted as hypothetical proteins (Figures 5 and 6).

**Figure 5.**
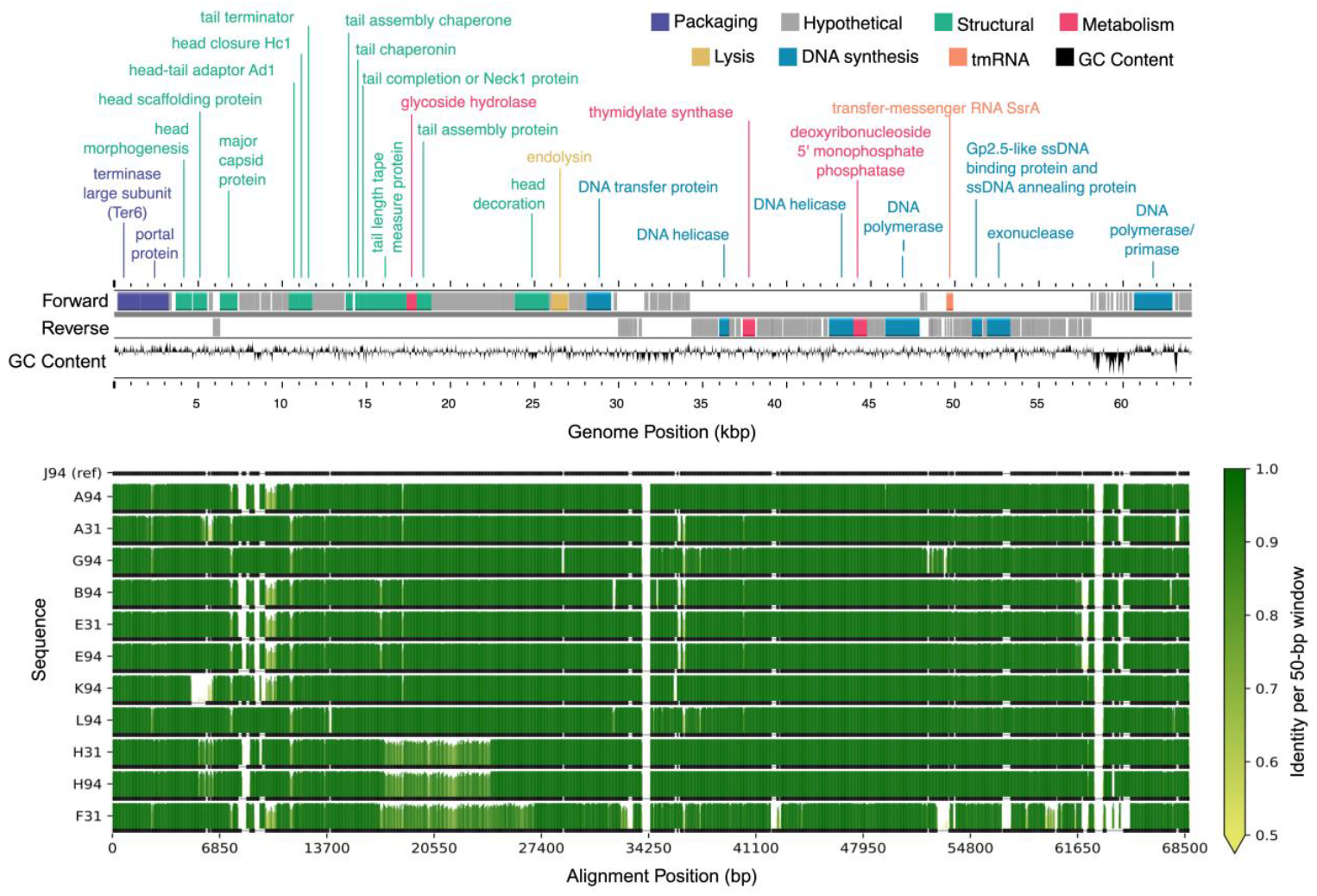
Genome map and alignment of 12 lytic phages isolated from Delaware soils against *Bradyrhizobium elkanii*. The top panel shows the genome map of representative phage J94 with forward and reverse strand gene annotations colored by predicted function and GC skew. The bottom panel displays pairwise genome alignments, with sequence identity relative to J94 (reference) across 50 bp windows shown as vertical bars. Horizontal black line below each identity plot indicates regions that are present (thick) versus absent (thin) for a given genome, accommodating insertions or deletions relative to all other sequences (including J94 reference).

**Figure 6.**
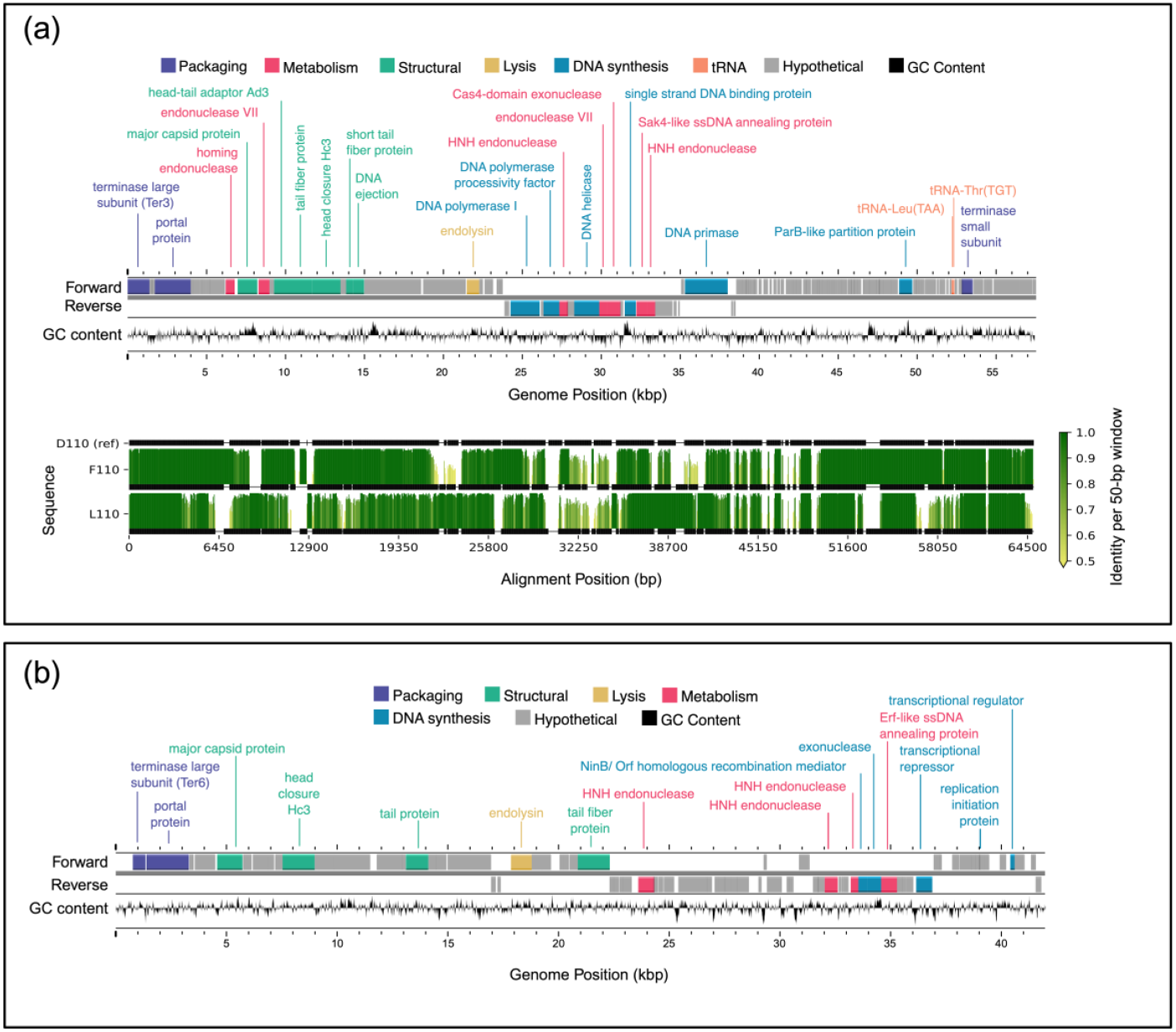
Genome maps and alignment of lytic phages isolated from Delaware soils against *Bradyrhizobium diazoefficiens*. (a) Genome map and alignment of three phages isolated against *B. diazoefficiens* which grouped together based on ANI. The top panel shows the genome map of representative phage D110 with forward and reverse strand gene annotations colored by predicted function and a GC skew plot. The bottom panel displays pairwise genome alignments to D110, with sequence identity across 50 bp windows shown as vertical bars. Horizontal black bars below each identity plot indicate sequenced portions of a given genome, and the thin horizontal black lines between bars indicate gaps introduced to accommodate insertions or deletions relative to the reference, including in D110 to match regions present in other genomes. (b) Genome map of phage J110, which formed a distinct group with no closely related genomes based on ANI. The genome map includes forward, and reverse strand gene annotations colored by predicted function, along with a GC skew plot. No alignment is shown due to lack of genomic similarity to the other phage isolates.

All 16 genomes contained an annotated large subunit terminase (TerL) gene (Figures 5 and 6). Pfam domain analysis of the TerL N-terminus showed that the 12 *B. elkanii* phages and *B. diazoefficiens* phage J110 encoded a terminase-6 domain, like that found in phage T7 [43]. In contrast, the three remaining *B. diazoefficiens* phages harbored a terminase-3 domain, as seen in phages such as *Shigella* phage Sf6 [43].Additionally, all except for J110 had an annotated major capsid protein and DNA polymerase I. Analysis of the *polA* 762 position (*E. coli* numbering), a key catalytic enzyme linked to processivity speed [46, 84-87], revealed the *B. elkanii* and *B. diazoefficiens* phages carry a leucine (Leu762) *polA* and a phenylalanine (Phe762) *polA*, respectively. All 16 phages encoded tail related genes, consistent with the siphophage and podophage morphology observed by TEM. Additionally, the *B. elkanii* phages carried a tmRNA gene, while the *B. diazoefficiens* phages (excluding J110) carried tRNA-Leu and tRNA-Thr genes.

### 3.7 Comparative Genome Alignments

Comparative genome alignments assessed nucleotide-level similarities and differences within the ANI-determined phage species (Figures 5 and 6). The longest genome from each group was selected as the reference sequence for alignment, specifically J94 and D110 for the *B. elkanii* and *B. diazoefficiens* phages, respectively; J110 was excluded from the *B. diazoefficiens* group as it shared less than 80% identity with the other phages and did not cluster with any group based on ANI. Among the *B. elkanii* phages, most nucleotide differences occurred within structural and tail component genes, while genes required for successful phage replication and packaging such as DNA polymerase I and *TerL* were more conserved. Notably, F31 exhibited greater dissimilarity across its entire genome compared to the other *B. elkanii* phages. F110 and L110 had less overall homology to the reference genome (D110) when compared to the *B. elkanii* phage group.

Supplemental genome comparisons were performed using Clinker and Anvi’o software. The Clinker alignment, like the ANI results, divided the 16 phages into three groups, with J110 representing a unique group (Figure S3). The alignments showed a higher degree of homology across the *B. elkanii* phages than for the *B. diazoefficiens* phages. A single hypothetical gene, located from approximately 14,900 – 16,900bp (Figure 6b), in the J110 genome shared amino acid and nucleotide-level homology with the other *B. diazoefficiens* phages (Figures S3 and S4). Structural prediction of this shared gene in J110 using AlphaFold, followed by Foldseek structural homology searches, indicated structural similarity (TM-scores >0.5) to tail-associated proteins such as tail fibers and tail spike proteins. In contrast, structural searches of the homolog in the other USDA110 phages returned top hits annotated as uncharacterized proteins, with no functional annotation.

The Anvi’o pangenome analysis showed a similar trend to Clinker, with gene clusters grouping with the ANI-determined phage groups (Figure S4). The pangenome identified 90 core gene clusters that were shared across all the *B. elkanii* phage genomes, and 82 that were shared among the *B. diazoefficiens* phages (excluding J110). There was one gene cluster, annotated with an unknown function, that was shared between J110 and the other three *B. diazoefficiens* phages; this was the only gene cluster that was found in more than one ANI-determined species (ANI ≥ 95%). Excluding phage J110, Anvi’o also identified 34 gene clusters, annotated as hypothetical proteins, that were unique, each found in only one of the following genomes: D110, F110, L110, E31, F31, and K94. J110 was excluded from this count because its entire genome (except for the one previously mentioned gene) was unique compared to the other 15 phages.

### 3.8 Geographical Analysis

Mantel tests were performed on three subsets of the ANI results (all 16 phages, only the *B. elkanii* phages, and only the *B. diazoefficiens* phages) assessing whether differences in nucleotide identity were associated geographic distribution of phage isolation sites. No significant correlation was detected between geographic distance and genomic similarity across all 16 phages, or within the *B. elkanii* and *B. diazoefficiens* subsets, suggesting broad genetic similarity across the geographic distribution of these phages.

## 4. Discussion

The isolation and characterization of 16 virulent *Bradyrhizobium* phages from Delaware soybean farms expands our understanding of soil viral diversity and functional potential in agricultural soils. While phages are known to regulate bacterial populations through lysis, horizontal gene transfer, and nutrient cycling [15,16], those infecting soybean-nodulating *Bradyrhizobium spp*. have remained largely uncharacterized. To our knowledge, this work constitutes the largest addition of *Bradyrhizobium* phage genomes to date, significantly expanding the genomic resources available for understanding agriculturally relevant soil phages.

### 4.1 Phage Isolation and Host Range

Phages were enriched from 10 Delaware agricultural soils against 10 host strains. Only three USDA strains yielded phages whereas no isolates were obtained from five native Delaware strains, two additional USDA strains, or four *B. japonicum* strains (Figure 1). The lack of phages isolated against native strains may reflect evolved immunity due to selective pressure by phage [59]. Six soil samples yielded phages infecting multiple host strains, with three of these yielding phages capable of infecting strains from different *Bradyrhizobium* species, suggesting diverse host and phage communities [34,60]. There are an estimated ∼10^9^ phages per gram of agricultural soil in Delaware [23,61]. Since only one plaque was isolated per soil–host combination, and many phages fail to produce visible plaques [61,62], the 16 isolates likely represent a fraction of *Bradyrhizobium* phage diversity. Although we did not characterize bradyrhizobia populations in our particular soils, *B. elkanii* is reportedly the most common species in the southeastern and mid-Atlantic U.S., particularly those strains serologically related to USDA 94 [63,64]. We speculate that the predominance of *B. elkanii* phages among our isolates may reflect “kill-the-winner” dynamics, where competitive host strains are preferentially targeted by phages [65, 66]. However, an expanded host range analysis and additional phage isolations are needed to fully support this claim.

Host range analysis showed species-specific infection patterns, with phages infecting only *Bradyrhizobium* strains of the same species against which they were isolated (Figure 3). All phages were able to infect multiple strains within their isolation host species, suggesting shared phage receptors among the susceptible strains and emphasizing the importance of bacterial taxonomy in phage host range [67]. Phages isolated against *B. diazoefficiens* lysed USDA 110 and S13E but not USDA 122, whereas *B. elkanii* phages lysed S01C, USDA 31, USDA 76, and USDA 94 regardless of whether the host was a Delaware isolate or USDA reference. Some *B. elkanii* strains (USDA 130, S05J) resisted all phages despite close ITS similarity to susceptible strains (USDA 31) [34], highlighting complex phage–host compatibility. While these results provide insight into the host specificity and infection range of the 16 isolated phages, it is important to recognize that infection patterns observed under laboratory conditions may not necessarily reflect what occurs in the natural soil environment [68]. Further investigation of phage-resistant strains could reveal mechanisms of immunity and identify highly effective bradyrhizobia symbionts for soybean inoculants to enhance nitrogen fixation [22].

### 4.2 Morphology

Morphological characterization using TEM divided the 16 phages into two groups dependent on the species of their isolation host. The *B. elkanii* phages all possessed long, non-contractile tails characteristic of siphophages and prolate capsids, whereas the phages isolated against *B. diazoefficiens* had icosahedral capsids and short podophage-like tails [33] (Figure 2). Previously described virulent phages infecting *Bradyrhizobium* have been classified within the podophage and siphophage morphological groups, with at least one study reporting both morphotypes capable of infecting *B. japonicum* strains [18,22]. While we did not isolate phages infecting *B. japonicum*, the phages recovered against *B. elkanii* and *B. diazoefficiens* exhibit morphological features consistent with those previously reported for *Bradyrhizobium*-infecting phages [22,25].

### 4.3 Genome Composition

Previous studies have reported inverse relationships between genome size and G+C content, as well as linear correlations between phage and host G+C content [69]. Our *B. elkanii* phages possessed relatively large genomes with G+C contents substantially higher than those of the *B. diazoefficiens* phages, despite their hosts having similar genomic G+C values [25]. Elevated G+C content is often linked to increased DNA stability and is particularly enriched in structural genes [70]. Consistent with this, we observed higher G+C levels in major capsid and head–tail complex proteins.

Phages tend to be more A+T rich than their hosts, which can make translation difficult if the phage relies entirely on the host’s codon pools [71,72]. All the phages isolated against USDA 110, excluding J110, carry at least two annotated tRNA genes (Figure 6) which are A+T biased. These phage-encoded tRNAs likely supplement host tRNAs, ensuring that the A+T-rich codons favored by the phage can be recognized and synthesized by the ribosome. Even though USDA 110 carries the same tRNAs in its genome, the phage-encoded copies may accelerate phage protein synthesis and reduce dependency on the host’s translational machinery, representing a potential adaptive strategy to optimize infection and phage replication [71].

While the *B. elkanii* phages did not encode tRNAs in their genomes, they did encode transfer-messenger RNAs (tmRNAs). tmRNAs, combining properties of both tRNA and mRNA, play a key role in rescuing stalled ribosomes and tagging incomplete polypeptides for degradation [73]. By encoding their own tmRNAs, these phages may enhance translation efficiency, particularly under conditions where stalled host ribosomes are limiting the phage replication process. The presence of tmRNAs in phage genomes is an example of strategies phages may use to manipulate their hosts’ machinery to optimize their replication.

### 4.4 Comparative Genomics

According to ICTV guidelines, phages sharing ≥95% ANI are classified as the same species [27]. Applying this framework, the 16 phages reported here represent three distinct species, highlighting the genomic diversity within *Bradyrhizobium* phages. (Figure 4). Notably, the *B. elkanii* phages exhibited exceptionally high ANI values, often exceeding 99.5%, which suggests the presence of subspecies-level groupings, or genomovars [30]. Given that many of our *B. elkanii* phages were isolated from sites separated by up to 70 km and yet still clustered within the same genomovar indicates a broad geographic distribution of highly similar phage populations within this region or strong selective pressures maintaining genomic conservation. Although F110, D110, and L110 did share a sufficiently high ANI to be considered members of the same species, the values were insufficient to further group them into genomovars.

Both the whole genome alignments (Figures 5 and 6) and Clinker visualizations (Figure S3) confirmed the ANI-based clustering of the 16 phages into three distinct groups, reinforcing the robustness of ANI as a metric for phage species delineation. The consistent separation of J110 from the other phages, along with its lack of genomic similarity aside from a single conserved hypothetical gene cluster shared with the other *B. diazoefficiens* phages (Figure S3), indicates it represents a distinct species or lineage from all of the other bradyrhizobia phages isolated in this study. Although J110 was highly divergent at the genomic level, its host range and morphology were the same as the other *B. diazoefficiens* phages (D110, F110, L110). This suggests that, despite being genetically distinct, J110 may have independently evolved or acquired infection strategies like those of the other *B. diazoefficiens* phages, highlighting possible convergent evolution within the group.

Among the *B. elkanii* phages, nucleotide-level variation appeared to be concentrated in genes related to structural and tail proteins, which are known to influence host specificity [74]. F31 exhibited more widespread dissimilarity across its genome, suggesting it may have undergone recombination or horizontal gene transfer events making its genome slightly more diverse. While F31 was the most divergent from the other *B. elkanii* phages, it must be reiterated that it still shared above 97% ANI and 85% coverage with these phages. The Anvi’o pangenome analysis also revealed a strong separation of gene clusters between the *B. elkanii* and *B. diazoefficiens* phage groups (Figure S4). This reinforces the idea that phage evolution is tightly linked to host phylogeny [75]. The identification of 34 unique gene clusters and a fully unique genome in J110 (aside from one conserved gene) suggests extensive phage diversification within the 16 *Bradyrhizobium* phages. These unique genes, all annotated as hypothetical proteins, represent candidates for future functional characterization. The presence of 90 core genes in the *B. elkanii* phages and 82 in the *B. diazoefficiens* group suggests strong conservation of essential functional elements within each phage group. The core genes included phage structural and replication proteins required for successful infection of their given host species.

### 4.5 Conserved Genes and Genetic Diversity

Approximately 24% of genes in each sequenced phage genome were annotated with a known function using Pharokka (Table S2). Similarly low annotation percentages are common in the literature and reflect the substantial genetic novelty found in phages [76]. This genetic diversity is driven largely by high rates of gene exchange and host-specific adaptations [77]. Among the annotated genes, those encoding the terminase large subunit (*TerL*), major capsid protein, and DNA polymerase I were the most conserved, with all 16 genomes containing a *TerL* gene and all but J110 possessing an annotated major capsid protein and DNA polymerase(Figures 5 and 6). These three genes are essential for phage structure, genome replication, and DNA packaging during lytic infection [78,79]. Within the 12 *B. elkanii* phages, tail gene regions exhibited the highest sequence variability, particularly in strain F31, which showed the most divergence at the nucleotide level (Figure 5). Although F31 infected the same strains as other *B. elkanii* phages, it only partially lysed USDA 94, whereas complete lysis was observed with the other isolates. Thus, the nucleotide variation observed in the F31 tail genes may have influenced its infection efficiency [74,80].

The genomic divergence of J110 from the other USDA110 phages, despite its shared host range, raises questions about which specific genomic features may contribute to host recognition in these phages. The only shared gene among the USDA110 phages was initially annotated as hypothetical. Structural homology search using Foldseek for the J110 allele produced top hits to tail-associated proteins, whereas the corresponding genes in the other USDA110 phages lacked strong matches to functionally annotated genes. Although sequence similarity analysis by Clinker (Figure S3) confirmed similarity among these genes in the USDA110 phages, they share only ∼25% identity and ∼35% similarity at the amino acid level. While the predicted tail-associated function in J110 suggests that this shared gene could contribute to host recognition [81], the ∼25% sequence identity between J110 and the other USDA110 phages falls within the “twilight zone” of protein homology [80], preventing a definitive functional annotation. This finding illustrates how uncharacterized proteins may have important structural or ecological functions [81], underscoring the need for further investigation and potential experimental validation.

### 4.6 Functional Insights into terL and polA Genes

The *TerL* gene plays a central role in DNA packaging, and its protein family (Pfam) domain can predict packaging mechanisms such as 3′ cohesive ends, 5′ cohesive ends, or headful packaging [43,44,83]. Based on the *TerL* Pfam annotations, the phages grouped into two distinct terminase families: the *B. elkanii* phages and J110 encoded terminase-6, while the remaining *B. diazoefficiens* phages encoded terminase-3 domains. Despite the differences, both domains are associated with headful packaging [43], a mechanism in which DNA packaging is driven by the volume and capacity of the capsid rather than sequence-specific termination sites. As a result, headful packaging can facilitate generalized transduction by incorporating host DNA into phage capsids, thereby promoting horizontal gene transfer [83].

Family A DNA polymerase, encoded by the *polA* gene, is the primary enzyme involved in phage genome replication and is found in approximately 25% of dsDNA phages [84]. The identity of the residue at the 762 position (*E. coli* numbering) has been shown to affect the processivity and fidelity of the enzyme [46,85–88]. Experimental mutagenesis of residue 762 in *E. coli polA* has shown that amino acid identity at this position alters enzymatic fidelity and processivity, with phenylalanine (Phe762, wild type) and tyrosine (Tyr762) substitutions associated with higher processivity rates [85], and leucine (Leu762) resulting in a slower but more accurate polymerase [86]. Genome-to-phenome connections have been made for phage *polA* sequences, suggesting the identity of the residue at the 762 position can be indicative of phage lifestyle. Previously reported evidence supports the hypothesis that the higher processivity of the Phe762 and Tyr762 variants is favorable to virulent phages, whereas the slower Leu762 variant is associated with temperate phages [89,90]. Analysis of the *polA* genes found in the *Bradyrhizobium* phages described in this study showed that the phages isolated against a *B. elkanii* host have a leucine in the 762 position (Leu762), whereas those isolated against a *B. diazoefficiens* host have a phenylalanine (Phe762). The finding of a Phe762 *polA* in the *B. diazoefficiens* phages supports prior evidence that this variant is favorable for a lytic lifestyle that may require rapid DNA synthesis for viral replication. However, the *B. elkanii* phages possessed a Leu762 variant typically associated with a temperate lifestyle, which diverges from this genome-to-phenome connection due to current evidence that these phages are lytic. A potential explanation for the presence of Leu762 *polA* in the virulent *B. elkanii* phages is the relatively long doubling time of *Bradyrhizobium* in laboratory culture compared to *E. coli*, which divides approximately 28-fold faster [43,91]. The slower host growth rate may reduce pressure on the phage to replicate rapidly, making the advantage of a faster polymerase less critical for successful phage replication. Additionally, although these phages exhibited lytic behavior under the given laboratory conditions tested in this study, it is possible that they are capable of lysogeny under different conditions or with other hosts [92]. The partial lysis observed in some of the *B. elkanii* phages during host range assays (Figure 3) is also suggestive of lysogeny, as lysogens often produce turbid plaques [33]. While it may be tempting to interpret the presence of Leu762 and partial lysis as evidence for lysogeny in the *B. elkanii* phages, alternative explanations cannot be ruled out. Notably, no hallmark genes of lysogeny were annotated in these genomes [93]. However, because a large proportion of their genes remain hypothetical, we cannot conclude that lysogeny-associated functions are completely absent, as some of these uncharacterized genes may encode integration-related functions.

### 4.7 Geographic Influence on Phage Diversity

One of the more remarkable findings of our study is the lack of a significant geographical effect on phage diversity, particularly among the *B. elkanii* phage isolates. The 12 phages isolated against this host species, despite being obtained from farms up to ∼70 km distant and enriched against two host strains, all shared ANI values >97% and similar genomic content and synteny. This high genomic similarity across geographical scales suggests the influence of host-driven genome conservation rather than geographic diversification. It is important to recall that these phages were isolated from soybean farms, which are highly managed environments strongly shaped by farming practices [94]. Additional work examining phages from unmanaged soils or increasing the number of sampling sites are necessary before drawing broader conclusions about the predominance of these phages in Delaware soils and the ecological roles they may play.

### 4.8 Conclusion

The findings of this study highlight the diversity, host-specificity, and conservation among *Bradyrhizobium*-infecting phages present in Delaware agricultural soils. A remarkable finding is that, despite being isolated from soils up to 70 km apart and the previously reported diversity of soybean bradyrhizobia in Delaware [63,64], phages isolated against *B. elkanii* exhibited high genomic similarity, with many phages in this group sharing over 99% ANI. Conversely, when phages isolated against *B. diazoefficiens* are also considered, two additional phage species were identified, including one (J110) with no shared genomic content aside from a single homologous gene, thus highlighting the complex nature of phage evolution. The results of the phenotypic analysis, as well as comparative genomics, support the idea that phage evolution is linked to host phylogeny. Additionally, the numerous hypothetical and unique genes identified in this study provide targets for future research into the molecular mechanisms of phage infection, adaptation, and evolution in soybean-*Bradyrhizobium* systems.

## Supplementary Materials

The following supporting information can be downloaded at: www.mdpi.com/xxx/s1. Table A1: Means and corresponding standard deviations of morphological characteristics for 16 virulent *Bradyrhizobium* phages isolated from Delaware soils. Table A2: Genomic features of 16 virulent *Bradyrhizobium* phages isolated from Delaware soils. Genome length in base pairs, percent GC content, total number of genes, and the number of functionally annotated genes are shown. Figure A1: Boxplots of capsid dimensions and estimated capsid volumes of the 16 virulent bacteriophages isolated against *Bradyrhizobium elkanii* and *B. diazoefficiens*. Figure A2: Representative host range spot assay results showing the varying levels of lytic activity by *Bradyrhizobium* phages. Figure A3: Clinker alignment of the whole genomes of 16 lytic *Bradyrhizobium* bacteriophages isolated from Delaware soils showing the separation of the phages at the protein and nucleotide level, congruent with ANI species groupings. Figure A4: Pangenome display of 16 phages virulent on soybean *Bradyrhizobium* spp. isolated from Delaware soils.

## Author Contributions

Conceptualization, E.A.M., J.J.F., K.E.W., S.W.P., and B.D.F.; methodology, E.A.M., J.J.F., K.E.W., S.W.P., and B.D.F.; formal analysis, E.A.M.; investigation, E.A.M., S.C.T.; resources, J.J.F.; writing—original draft preparation, E.A.M..; writing—review and editing, E.A.M., J.J.F., K.E.W., S.W.P., S.C.T., and B.D.F.; visualization, E.A.M., J.J.F. and S.W.P.; supervision, J.J.F., K.E.W., S.W.P., and B.D.F.; project administration, B.D.F.; funding acquisition, K.E.W., S.W.P., J.J.F., and B.D.F. All authors have read and agreed to the published version of the manuscript.

## Institutional Review Board Statement

Not applicable

## Informed Consent Statement

Not applicable

## Data Availability Statement

The raw data supporting the conclusions of this article will be made available by the authors on request. DNA sequences of the phage isolates are available at NCBI under Bioproject PRJNAxxxxxx and GenBank accession numbers PX020967-PX020982.

## Acknowledgments

We thank Shannon Modla (University of Delaware Bio-Imaging Center) for TEM expertise and virion imaging. Student financial support for E.A.M was provided by the following University of Delaware-based sources: Fellowship from the Microbiology Graduate Program and Unidel Foundation, Graduate Scholars Award, Department of Plant and Soil Sciences teaching assistantship, Department of Biological Sciences teaching assistantship. Student financial support for S.C.T. was provided by the following University of Delaware-based sources: Environmental Science Department Internship Award, Undergraduate Research Summer Scholar Award, Delaware Environmental Institute (DENIN) Environmental Scholar Award, and Harward Munson Fellowship. Support from the University of Delaware Bioinformatics Data Science Core Facility (RRID:SCR_017696), the University of Delaware Sequencing and Genotyping. Center (RRID:SCR_012230), and use of the BIOMIX and BioStoRe compute resources was made possible through funding from Delaware INBRE (NIH P20GM103446), an NIH Shared Instrumentation Grant (NIH S10OD028725), the State of Delaware, and the Delaware Biotechnology Institute.

## Disclaimer/Publisher’s Note

The statements, opinions and data contained in all publications are solely those of the individual author(s) and contributor(s) and not of MDPI and/or the editor(s). MDPI and/or the editor(s) disclaim responsibility for any injury to people or property resulting from any ideas, methods, instructions or products referred to in the content.

## References

1. Soybeans | USDA Foreign Agricultural Service Available online: https://www.fas.usda.gov/data/production/commodity/2222000 (accessed on 30 May 2025).

2. Sugiyama, A.; Ueda, Y.; Takase, H.; Yazaki, K. Do Soybeans Select Specific Species of Bradyrhizobium during Growth? Commun Integr Biol 2015, 8, doi:10.4161/19420889.2014.992734.

3. Messina, M. Perspective: Soybeans Can Help Address the Caloric and Protein Needs of a Growing Global Population. Front Nutr 2022, 9, doi:10.3389/FNUT.2022.909464.

4. Shober, A.L.; Taylor, R. Nitrogen Management for Soybeans | Cooperative Extension | University of Delaware Available online: https://www.udel.edu/academics/colleges/canr/cooperative-extension/fact-sheets/nitrogen-management-soybeans/ (accessed on 30 May 2025).

5. Tamagno, S.; Sadras, V.O.; Haegele, J.W.; Armstrong, P.R.; Ciampitti, I.A. Interplay between Nitrogen Fertilizer and Biological Nitrogen Fixation in Soybean: Implications on Seed Yield and Biomass Allocation. Scientific Reports 2018 8:1 2018, 8, 1–11, doi:10.1038/s41598-018-35672-1.

6. Menegat, S.; Ledo, A.; Tirado, R. Greenhouse Gas Emissions from Global Production and Use of Nitrogen Synthetic Fertilisers in Agriculture. Sci Rep 2022, 12, doi:10.1038/S41598-022-18773-W.

7. Norton, J.; Ouyang, Y. Controls and Adaptive Management of Nitrification in Agricultural Soils. Front Microbiol 2019, 10, 449199, doi:10.3389/FMICB.2019.01931/XML/NLM.

8. Wang, J.; Liu, X.; Beusen, A.H.W.; Middelburg, J.J. Surface-Water Nitrate Exposure to World Populations Has Expanded and Intensified during 1970-2010. Environ Sci Technol 2023, 57, 19395–19406, doi:10.1021/ACS.EST.3C06150/SUPPL_FILE/ES3C06150_SI_001.PDF.

9. Paerl, H.W. Coastal Eutrophication and Harmful Algal Blooms: Importance of Atmospheric Deposition and Groundwater as “New” Nitrogen and Other Nutrient Sources. Limnol Oceanogr 1997, 42, 1154–1165, doi:10.4319/LO.1997.42.5_PART_2.1154.

10. Tian, H.; Xu, R.; Canadell, J.G.; Thompson, R.L.; Winiwarter, W.; Suntharalingam, P.; Davidson, E.A.; Ciais, P.; Jackson, R.B.; Janssens-Maenhout, G.; et al. A Comprehensive Quantification of Global Nitrous Oxide Sources and Sinks. Nature 2020 586:7828 2020, 586, 248–256, doi:10.1038/s41586-020-2780-0.

11. Fuhrmann, J.J.; Vasilas, B.L. Field Response of the Glycine-Bradyrhizobium Symbiosis to Modified Early-Nodule Occupancy. Soil Biol Biochem 1993, 25, 1203–1209, doi:10.1016/0038-0717(93)90216-X.

12. Zimmer, S.; Messmer, M.; Haase, T.; Piepho, H.-P.; Mindermann, A.; Schulz, H.; Habekuß, A.; Ordon, F.; Wilbois, K.-P.; Heß, J. Effects of Soybean Variety and Bradyrhizobium Strains on Yield, Protein Content and Biological Nitrogen Fixation under Cool Growing Conditions in Germany. Europ. J. Agronomy 2016, 72, 38–46, doi:10.1016/j.eja.2015.09.008.

13. Mushegian, A.R. Are There 1031virus Particles on Earth, or More, or Fewer? J Bacteriol 2020, 202, doi:10.1128/JB.00052-20.

14. Hendrix, R.W.; Smith, M.C.M.; Burns, R.N.; Ford, M.E.; Hatfull, G.F. Evolutionary Relationships among Diverse Bacteriophages and Prophages: All the World’s a Phage. Proc Natl Acad Sci U S A 1999, 96, 2192–2197, doi:10.1073/pnas.96.5.2192.

15. Domingo, E. Quasispecies Dynamics in Disease Prevention and Control. Virus as Populations 2016, 263–297, doi:10.1016/B978-0-12-800837-9.00008-3.

16. Williamson, K.E.; Fuhrmann, J.J.; Wommack, K.E.; Radosevich, M. Viruses in Soil Ecosystems: An Unknown Quantity within an Unexplored Territory. Annu Rev Virol 2017, 4, 201–219, doi:10.1146/ANNUREV-VIROLOGY-101416-041639.

17. Ali, F.S.; Loynachan, T.E.; Hammad, A.M.M.; Aharchi, Y. Polyvirulent Rhizobiophage from a Soybean Rhizosphere Soil. Soil Biol Biochem 1998, 30, 2171–2175, doi:10.1016/S0038-0717(98)00048-0.

18. Appunu, C.; Dhar, B. Morphology and General Characteristics of Lytic Phages Infective on Strains of Bradyrhizobium japonicum. Curr Microbiol 2008, 56, 21–27, doi:10.1007/s00284-007-9031-6.

19. Appunu, C.; Dhar, B. Isolation and Symbiotic Characteristics of Two Tn5-Derived Phage-Resistant Bradyrhizobium japonicum Strains That Nodulate Soybean. Curr Microbiol 2008, 57, 212–217, doi:10.1007/S00284-008-9176-Y.

20. Hashem, F.M.; Angle, J.S. Rhizobiophage Effects on Bradyrhizobium japonicum, Nodulation and Soybean Growth. Soil Biol Biochem 1988, 20, 69–73, doi:10.1016/0038-0717(88)90128-9.

21. Hashem, F.M.; Angle, J.S.; Ristiano, P.A. Isolation and Characterization of Rhizobiophages Specific for Bradyrhizobium japonicum USDA 117. Can J Microbiol 1986, 32, 326–329, doi:10.1139/M86-064.

22. Msimbira, L.A.; Jaiswal, S.K.; Dakora, F.D. Identification and Characterization of Phages Parasitic on Bradyrhizobia Nodulating Groundnut (Arachis Hypogaea L.) in South Africa. Applied Soil Ecology 2016, 108, 334–340, doi:10.1016/J.APSOIL.2016.09.010.

23. Shahaby, A.F.; Alharthi, A.A.; El-Tarras, A.E. Characterization of Rhizobiophages Specific for Rhizobium Sp. Sinorhizobum Sp., and Bradyrhizobium Sp. Int J Curr Microbiol Appl Sci 2014, 3, 155–171.

24. Kowalski, M.; Ham, G.E.; Frederick, L.R.; Anderson, I.C. Relationship between Strains of Rhizobium japonicum and Their Bacteriophages from Soil and Nodules of Field-Grown Soybeans. Soil Sci 1974, 118, 221–228.

25. Richards, V.A.; Ferrell, B.D.; Polson, S.W.; Wommack, K.E.; Fuhrmann, J.J. Soybean Bradyrhizobium Spp. Spontaneously Produce Abundant and Diverse Temperate Phages in Culture. Viruses 2024, 16, 1750, doi:10.3390/V16111750/S1.

26. Oliveira, H.; Domingues, R.; Evans, B.; Sutton, J.M.; Adriaenssens, E.M.; Turner, D. Genomic Diversity of Bacteriophages Infecting the Genus Acinetobacter. Viruses 2022, 14, doi:10.3390/V14020181.

27. Adriaenssens, E.M.; Brister, J.R. How to Name and Classify Your Phage: An Informal Guide. 2017, doi:10.1101/111526.

28. Turner, D.; Kropinski, A.M.; Adriaenssens, E.M. A Roadmap for Genome-Based Phage Taxonomy. Viruses 2021, 13, doi:10.3390/V13030506.

29. Ackermann, H.W. Bacteriophage Taxonomy. Microbiol Aust 2011, 32, 90, doi:10.1071/MA11090.

30. Aldeguer-Riquelme, B.; Conrad, R.E.; Antón, J.; Rossello-Mora, R.; Konstantinidis, K.T. A Natural ANI Gap That Can Define Intra-Species Units of Bacteriophages and Other Viruses. mBio 2024, 15, doi:10.1128/MBIO.01536-24/SUPPL_FILE/MBIO.01536-24-S0002.XLSX.

31. Chaudhari, H. V.; Inamdar, M.M.; Kondabagil, K. Scaling Relation between Genome Length and Particle Size of Viruses Provides Insights into Viral Life History. iScience 2021, 24, doi:10.1016/J.ISCI.2021.102452.

32. Dion, M.B.; Oechslin, F.; Moineau, S. Phage Diversity, Genomics and Phylogeny. Nat Rev Microbiol 2020, 18, 125–138.

33. Valencia-Toxqui, G.; Ramsey, J. How to Introduce a New Bacteriophage on the Block: A Short Guide to Phage Classification. 2024, doi:10.1128/jvi.01821-23.

34. Joglekar, P.; Mesa, C.P.; Richards, V.A.; Polson, S.W.; Wommack, K.E.; Fuhrmann, J.J. Polyphasic Analysis Reveals Correlation between Phenotypic and Genotypic Analysis in Soybean Bradyrhizobia (Bradyrhizobium Spp.). Syst Appl Microbiol 2020, 43, doi:10.1016/J.SYAPM.2020.126073.

35. Schindelin, J.; Arganda-Carreras, I.; Frise, E.; Kaynig, V.; Longair, M.; Pietzsch, T.; Preibisch, S.; Rueden, C.; Saalfeld, S.; Schmid, B.; et al. Fiji: An Open-Source Platform for Biological-Image Analysis. Nat Methods 2012, 9, 676–682, doi:10.1038/NMETH.2019.

36. Cui, J.; Schlub, T.E.; Holmes, E.C. An Allometric Relationship between the Genome Length and Virion Volume of Viruses. J Virol 2014, 88, 6403–6410, doi:10.1128/JVI.00362-14.

37. Wickham, H. Ggplot2; Use R!; Springer International Publishing: Cham, 2016; ISBN 978-3-319-24275-0.

38. Klindworth, A.; Pruesse, E.; Schweer, T.; Peplies, J.; Quast, C.; Horn, M.; Glöckner, F.O. Evaluation of General 16S Ribosomal RNA Gene PCR Primers for Classical and Next-Generation Sequencing-Based Diversity Studies. Nucleic Acids Res 2013, 41, doi:10.1093/NAR/GKS808.

39. Bouras, G.; Nepal, R.; Houtak, G.; Psaltis, A.J.; Wormald, P.J.; Vreugde, S. Pharokka: A Fast Scalable Bacteriophage Annotation Tool. Bioinformatics 2023, 39, doi:10.1093/BIOINFORMATICS/BTAC776.

40. Hyatt, D.; Chen, G.L.; LoCascio, P.F.; Land, M.L.; Larimer, F.W.; Hauser, L.J. Prodigal: Prokaryotic Gene Recognition and Translation Initiation Site Identification. BMC Bioinformatics 2010, 11, 1–11, doi:10.1186/1471-2105-11-119/TABLES/5.

41. Merrill, B.D.; Ward, A.T.; Grose, J.H.; Hope, S. Software-Based Analysis of Bacteriophage Genomes, Physical Ends, and Packaging Strategies. BMC Genomics 2016, 17, doi:10.1186/s12864-016-3018-2.

42. Finn, R.D.; Bateman, A.; Clements, J.; Coggill, P.; Eberhardt, R.Y.; Eddy, S.R.; Heger, A.; Hetherington, K.; Holm, L.; Mistry, J.; et al. Pfam: The Protein Families Database. Nucleic Acids Res 2013, 42, D222, doi:10.1093/NAR/GKT1223.

43. Joglekar, P.; Ferrell, B.D.; Jarvis, T.; Haramoto, K.; Place, N.; Dums, J.T.; Polson, S.W.; Wommack, K.E.; Fuhrmann, J.J. Spontaneously Produced Lysogenic Phages Are an Important Component of the Soybean Bradyrhizobium Mobilome. mBio 2023, 14, doi:10.1128/MBIO.00295-23.

44. Duffy, C.; Feiss, M. The Large Subunit of Bacteriophage λ’s Terminase Plays a Role in DNA Translocation and Packaging Termination. J Mol Biol 2002, 316, 547–561, doi:10.1006/JMBI.2001.5368.

45. Casjens, S. Prophages and Bacterial Genomics: What Have We Learned so Far? Mol Microbiol 2003, 49, 277–300.

46. Keown, R.A.; Dums, J.T.; Brumm, P.J.; MacDonald, J.; Mead, D.A.; Ferrell, B.D.; Moore, R.M.; Harrison, A.O.; Polson, S.W.; Wommack, K.E. Novel Viral DNA Polymerases from Metagenomes Suggest Genomic Sources of Strand-Displacing Biochemical Phenotypes. Front Microbiol 2022, 13, 858366, doi:10.3389/FMICB.2022.858366/BIBTEX.

47. Abramson, J.; Adler, J.; Dunger, J.; Evans, R.; Green, T.; Pritzel, A.; Ronneberger, O.; Willmore, L.; Ballard, A.J.; Bambrick, J.; et al. Accurate Structure Prediction of Biomolecular Interactions with AlphaFold 3. Nature 2024 630:8016 2024, 630, 493–500, doi:10.1038/s41586-024-07487-w.

48. van Kempen, M.; Kim, S.S.; Tumescheit, C.; Mirdita, M.; Lee, J.; Gilchrist, C.L.M.; Söding, J.; Steinegger, M. Fast and Accurate Protein Structure Search with Foldseek. Nature Biotechnology 2023 42:2 2023, 42, 243–246, doi:10.1038/s41587-023-01773-0.

49. Millard, A.; Denise, R.; Lestido, M.; Thomas, M.T.; Webster, D.; Turner, D.; Sicheritz-Pontén, T. TaxMyPhage: Automated Taxonomy of dsDNA Phage Genomes at the Genus and Species Level. Phage (New Rochelle) 2025, 6, 5–11, doi:10.1089/PHAGE.2024.0050.

50. Simmonds, P.; Adriaenssens, E.M.; Lefkowitz, E.J.; Oksanen, H.M.; Siddell, S.G.; Zerbini, F.M.; Alfenas-Zerbini, P.; Aylward, F.O.; Dempsey, D.M.; Dutilh, B.E.; et al. Changes to Virus Taxonomy and the ICTV Statutes Ratified by the International Committee on Taxonomy of Viruses. Archives of Virology 2024 169:11 2024, 169, 1–12, doi:10.1007/S00705-024-06143-Y.

51. Jain, C.; Rodriguez-R, L.M.; Phillippy, A.M.; Konstantinidis, K.T.; Aluru, S. High Throughput ANI Analysis of 90K Prokaryotic Genomes Reveals Clear Species Boundaries. Nature Communications 2018 9:1 2018, 9, 1–8, doi:10.1038/s41467-018-07641-9.

52. Grant, J.R.; Enns, E.; Marinier, E.; Mandal, A.; Herman, E.K.; Chen, C.Y.; Graham, M.; Van Domselaar, G.; Stothard, P. Proksee: In-Depth Characterization and Visualization of Bacterial Genomes. Nucleic Acids Res 2023, 51, W484–W492, doi:10.1093/NAR/GKAD326.

53. Gilchrist, C.L.M.; Chooi, Y.H. clinker & clustermap.js: Automatic Generation of Gene Cluster Comparison Figures. Bioinformatics 2021, 37, 2473–2475, doi:10.1093/BIOINFORMATICS/BTAB007.

54. Delmont, T.O.; Eren, E.M. Linking Pangenomes and Metagenomes: The Prochlorococcus Metapangenome. PeerJ 2018, 2018, doi:10.7717/PEERJ.4320.

55. Eren, A.M.; Kiefl, E.; Shaiber, A.; Veseli, I.; Miller, S.E.; Schechter, M.S.; Fink, I.; Pan, J.N.; Yousef, M.; Fogarty, E.C.; et al. Community-Led, Integrated, Reproducible Multi-Omics with Anvi’o. Nat Microbiol 2021, 6, 3–6, doi:10.1038/S41564-020-00834-3.

56. Mantel, N. The Detection of Disease Clustering and a Generalized Regression Approach. Cancer Res 1967, 27, 209–220.

57. Pedregosa, F.; Varoquaux, G.; Gramfort, A.; Michel, V.; Thirion, B.; Grisel, O.; Blondel, M.; Prettenhofer, P.; Weiss, R.; Dubourg, V.; et al. Scikit-Learn: Machine Learning in Python. Journal of Machine Learning Research 2011, 12, 2825–2830.

58. Lopez Gonzalez-Nieto, P.; Gomez Flechoso, M.; Arribas Mocoroa, M.E.; Muñoz Martin, A.; Garcia Lorenzo, M.L.; Cabrera Gomez, G.; Alvarez Gomez, J.A.; Caso Fraile, A.; Orosco Dagan, J.M.; Merinero Palomares, R.; et al. Design and Development of a Virtual Laboratory in Python for the Teaching of Data Analysis and Mathematics in Geology: GeoPy. INTED2020 Proceedings 2020, 1, 2236–2242, doi:10.21125/INTED.2020.0687.

59. Gómez, P.; Bennie, J.; Gaston, K.J.; Buckling, A. The Impact of Resource Availability on Bacterial Resistance to Phages in Soil. PLoS One 2015, 10, e0123752, doi:10.1371/JOURNAL.PONE.0123752.

60. Fuhrmann, J. Symbiotic Effectiveness of Indigenous Soybean Bradyrhizobia as Related to Serological, Morphological, Rhizobitoxine, and Hydrogenase Phenotypes. Appl Environ Microbiol 1990, 56, 224–229, doi:10.1128/AEM.56.1.224-229.1990.

61. Williamson, K.E.; Wommack, K.E.; Radosevich, M. Sampling Natural Viral Communities from Soil for Culture-Independent Analyses. Appl Environ Microbiol 2003, 69, 6628–6633, doi:10.1128/AEM.69.11.6628-6633.2003.

62. Williamson, K.E.; Radosevich, M.; Wommack, K.E. Abundance and Diversity of Viruses in Six Delaware Soils. Appl Environ Microbiol 2005, 71, 3119–3125, doi:10.1128/AEM.71.6.3119-3125.2005.

63. Fuhrmann, J. Serological Distribution of Bradyrhizobium japonicum as Influenced by Soybean Cultivar and Sampling Location. Soil Biol Biochem 1989, 21, 1079–1081, doi:10.1016/0038-0717(89)90047-3.

64. Minamisawa, K.; Onodera, S.; Tanimura, Y.; Kobayashi, N.; Yuhashi, K.I.; Kubota, M. Preferential Nodulation of Glycine Max, Glycine Soja and Macroptilium Atropurpureum by Two Bradyrhizobium Species japonicum and elkanii. FEMS Microbiol Ecol 1997, 24, 49–56, doi:10.1016/S0168-6496(97)00043-3.

65. Korytowski, D.A.; Smith, H. Permanence and Stability of a Kill the Winner Model in Marine Ecology. Bull Math Biol 2017, 79, 995–1004, doi:10.1007/S11538-017-0265-6.

66. Winter, C., Bouvier, T., Weinbauer, M. G., & Thingstad, T. F. Trade-offs between competition and defense specialists among unicellular planktonic organisms: the “killing the winner” hypothesis revisited. Microbiology and molecular biology reviews : MMBR 2010, 74(1), 42–57. 10.1128/MMBR.00034-09

67. Holtappels, D.; Alfenas-Zerbini, P.; Koskella, B. Drivers and Consequences of Bacteriophage Host Range. FEMS Microbiol Rev 2023, 47, doi:10.1093/FEMSRE/FUAD038.

68. Hyman, P.; Abedon, S.T. Bacteriophage Host Range and Bacterial Resistance. Adv Appl Microbiol 2010, 70, 217–248, doi:10.1016/S0065-2164(10)70007-1.

69. Almpanis, A.; Swain, M.; Gatherer, D.; McEwan, N. Correlation between Bacterial G+C Content, Genome Size and the G+C Content of Associated Plasmids and Bacteriophages. Microb Genom 2018, 4, doi:10.1099/MGEN.0.000168.

70. Das, R.; Rahlff, J. Phage Genome Architecture and GC Content: Structural Genes and Where to Find Them. 2024, doi:10.1101/2024.06.05.597531.

71. Bailly-Bechet, M.; Vergassola, M.; Rocha, E. Causes for the Intriguing Presence of tRNAs in Phages. Genome Res 2007, 17, 1486, doi:10.1101/GR.6649807.

72. Rocha, P.C.E.; Danchin, A. Base Composition Bias Might Result from Competition for Metabolic Resources. TRENDS in Genetics 2002, 18.

73. Ivanova, N.; Lindell, M.; Pavlov, M.; Schiavone, L.H.; Wagner, E.G.H.; Ehrenberg, M. Structure Probing of tmRNA in Distinct Stages of Trans-Translation. RNA 2007, 13, 713–722, doi:10.1261/RNA.451507.

74. Taslem Mourosi, J.; Awe, A.; Guo, W.; Batra, H.; Ganesh, H.; Wu, X.; Zhu, J. Understanding Bacteriophage Tail Fiber Interaction with Host Surface Receptor: The Key “Blueprint” for Reprogramming Phage Host Range. Int J Mol Sci 2022, 23, 12146, doi:10.3390/IJMS232012146.

75. Stern, A.; Sorek, R. The Phage-Host Arms-Race: Shaping the Evolution of Microbes. Bioessays 2011, 33, 43, doi:10.1002/BIES.201000071.

76. Ryu, S. Grand Challenges in Phage Biology. Front Microbiol 2021, 12, doi:10.3389/FMICB.2021.715039.

77. Grose, J.H.; Casjens, S.R. Understanding the Enormous Diversity of Bacteriophages: The Tailed Phages That Infect the Bacterial Family Enterobacteriaceae. Virology 2014, 468–470, 421–443, doi:10.1016/J.VIROL.2014.08.024.

78. Wangchuk, J.; Chatterjee, A.; Patil, S.; Madugula, S.K.; Kondabagil, K. The Coevolution of Large and Small Terminases of Bacteriophages Is a Result of Purifying Selection Leading to Phenotypic Stabilization. Virology 2021, 564, 13–25, doi:10.1016/J.VIROL.2021.09.004.

79. Choi, K.H. Viral Polymerases. Adv Exp Med Biol 2012, 726, 267–304, doi:10.1007/978-1-4614-0980-9_12.

80. Rost, B. Twilight Zone of protein sequence alignments. Protein Engineering, Design and Selection 1999, 12(2), 85–94. 10.1093/protein/12.2.85

81. Nobrega, F.L.; Vlot, M.; de Jonge, P.A.; Dreesens, L.L.; Beaumont, H.J.E.; Lavigne, R.; Dutilh, B.E.; Brouns, S.J.J. Targeting Mechanisms of Tailed Bacteriophages. Nature Reviews Microbiology 2018 16:12 2018, 16, 760–773, doi:10.1038/s41579-018-0070-8.

82. Vincent, A.T. Bacterial Hypothetical Proteins May Be of Functional Interest. Frontiers in Bacteriology 2024, 3, doi:10.3389/fbrio.2024.1334712.

83. Casjens, S.; Weigele, P. DNA Packaging by Bacteriophage P22. In Madame Curie Bioscience Database [Internet]; Landes Bioscience, 2013.

84. Wommack, K.E.; Nasko, D.J.; Chopyk, J.; Sakowski, E.G. Counts and Sequences, Observations That Continue to Change Our Understanding of Viruses in Nature. Journal of Microbiology 2015, 53, 181–192, doi:10.1007/S12275-015-5068-6/METRICS.

85. Astatke, M.; Ng, K.; Grindley, N.D.F.; Joyce, C.M. A Single Side Chain Prevents Escherichia coli DNA Polymerase I (Klenow Fragment) from Incorporating Ribonucleotides. Biochemistry 1998, 95, 3402–3407.

86. Suzuki, M.; Yoshida, S.; Adman, E.T.; Blank, A.; Loeb, L.A.; Gottstein, J. Thermus Aquaticus DNA Polymerase I Mutants with Altered Fidelity. Interacting Mutations in the O-Helix. Journal of Biological Chemistry 2000, 275, 32728–32735, doi:10.1074/JBC.M000097200/ASSET/422B8ED4-EC58-4146-8EB0-9F4CFA109EE7/MAIN.ASSETS/GR2.JPG.

87. Tabor, S.; Richardson, C.C. A Single Residue in DNA Polymerases of the Escherichia coli DNA Polymerase I Family Is Critical for Distinguishing between Deoxy- and Dideoxyribonucleotides. PNAS 1995, 92, 6339–6343.

88. Tabor, S.; Richardson, C.C. DNA Sequence Analysis with a Modified Bacteriophage T7 DNA Polymerase (DNA Polymerase I/Reverse Transcriptase/Chain-Terminating Inhibitors/2’-Deoxyinosine 5’-Triphosphate/Processivity). Biochemistry 1987, 84, 4767–4771.

89. Nasko, D.J.; Chopyk, J.; Sakowski, E.G.; Ferrell, B.D.; Polson, S.W.; Wommack, K.E. Family A DNA Polymerase Phylogeny Uncovers Diversity and Replication Gene Organization in the Virioplankton. Front Microbiol 2018, 9, 425215, doi:10.3389/FMICB.2018.03053/BIBTEX.

90. Schmidt, H.F.; Sakowski, E.G.; Williamson, S.J.; Polson, S.W.; Wommack, K.E. Shotgun Metagenomics Indicates Novel Family A DNA Polymerases Predominate within Marine Virioplankton. ISME J 2014, 8, 103–114, doi:10.1038/ismej.2013.124.

91. Tuttle, A.R.; Trahan, N.D.; Son, M.S. Growth and Maintenance of Escherichia coli Laboratory Strains. Curr Protoc 2021, 1, e20, doi:10.1002/CPZ1.20.

92. Howard-Varona, C., Hargreaves, K. R., Abedon, S. T., & Sullivan, M. B. Lysogeny in nature: mechanisms, impact and ecology of temperate phages. The ISME journal 2017 11(7), 1511–1520. 10.1038/ismej.2017.16

93. Altamirano, F. L. G., & Barr, J. J. Screening for Lysogen Activity in Therapeutically Relevant Bacteriophages. Bio-protocol 2021, 11(8), e3997. 10.21769/BioProtoc.3997

94. Vargas Gil, S.; Meriles, J.; Conforto, C.; Basanta, M.; Radl, V.; Hagn, A.; Schloter, M.; March, G.J. Response of Soil Microbial Communities to Different Management Practices in Surface Soils of a Soybean Agroecosystem in Argentina. Eur J Soil Biol 2011, 47, 55–60, doi:10.1016/J.EJSOBI.2010.11.006.

