## Supplemental figure 1 for "Comparative Analysis Reveals Host Species-Dependent Diversity Among 16 Virulent Bacteriophages Isolated Against Soybean *Bradyrhizobium* spp"

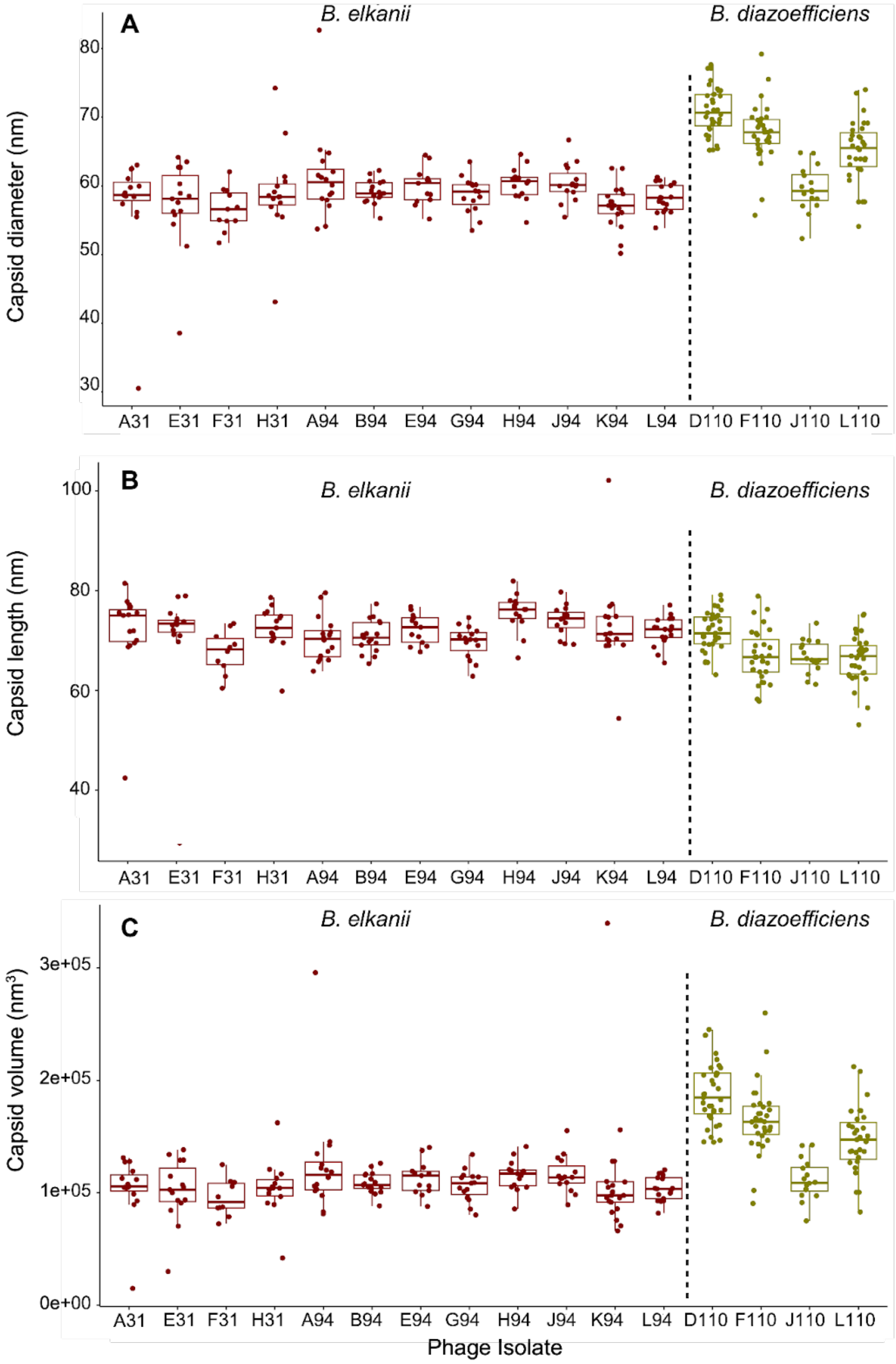


**Figure S1**. Boxplots of capsid dimensions and estimated capsid volumes of the 16 virulent bacteriophages isolated against *Bradyrhizobium elkanii* and *B. diazoefficiens*. Phage isolates are listed on the x-axis, with the capsid dimensions and volumes shown on the y-axis: (A) capsid diameter, (B) capsid length, and (C) capsid volume. Individual box plots and data points are colored according to the species of *Bradyrhizobium* used for isolation: *B. elkanii*, red; *B. diazoefficiens*, green.
