## Supplemental figure 2 for "Comparative Analysis Reveals Host Species-Dependent Diversity Among 16 Virulent Bacteriophages Isolated Against Soybean *Bradyrhizobium* spp"

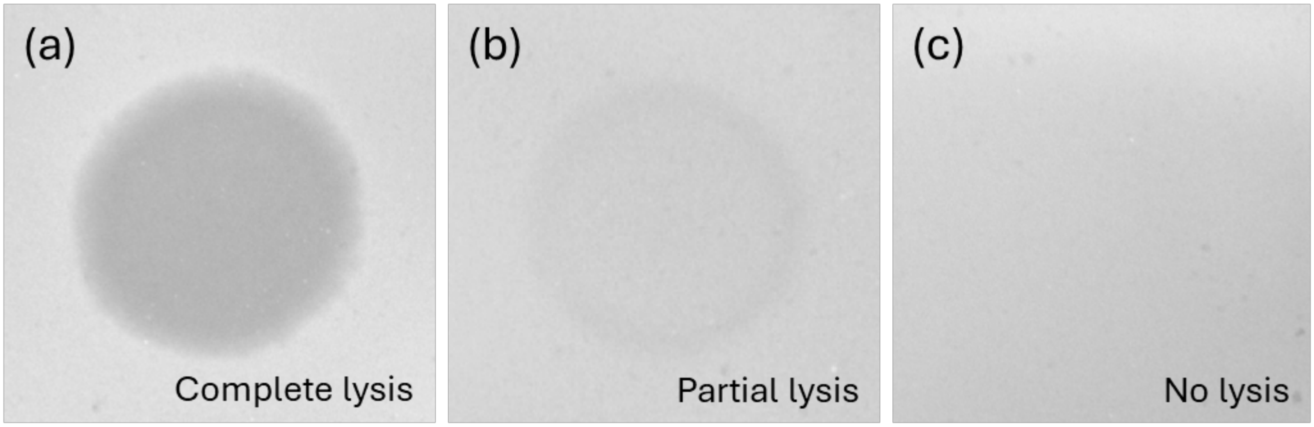


**Figure S2.** Representative host range spot assay results showing the varying levels of lytic activity by *Bradyrhizobium* phages. (a) Complete lysis: clear zone (b) Partial lysis: turbid zone (c) No lysis: absence of clearing.
