## Supplemental figure 3 for "Comparative Analysis Reveals Host Species-Dependent Diversity Among 16 Virulent Bacteriophages Isolated Against Soybean *Bradyrhizobium* spp"

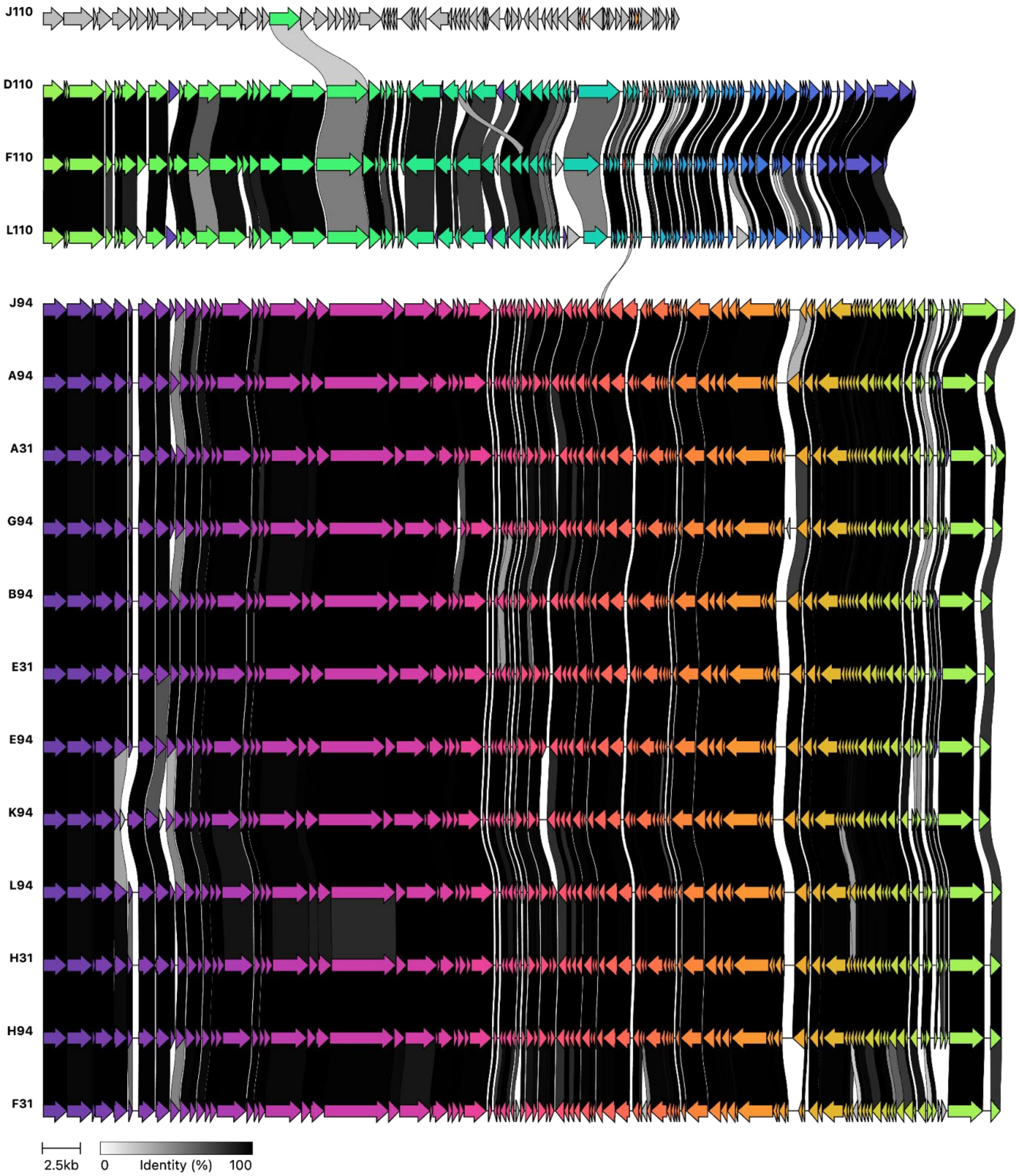


**Figure S3.** Clinker alignment of the complete genomes of 16 lytic *Bradyrhizobium* phages isolated from Delaware soils, showing protein and nucleotide level similarity consistent with ANI-based species groupings. Each row represents a phage genome, oriented from the large subunit terminase gene. Arrows denote individual genes, scaled by length and colored by Clinker-defined protein clusters; direction indicates strand orientation. Links between adjacent genomes represent nucleotide-level identity, shaded by percent identity.
