## Supplemental figure 4 for "Comparative Analysis Reveals Host Species-Dependent Diversity Among 16 Virulent Bacteriophages Isolated Against Soybean *Bradyrhizobium* spp"

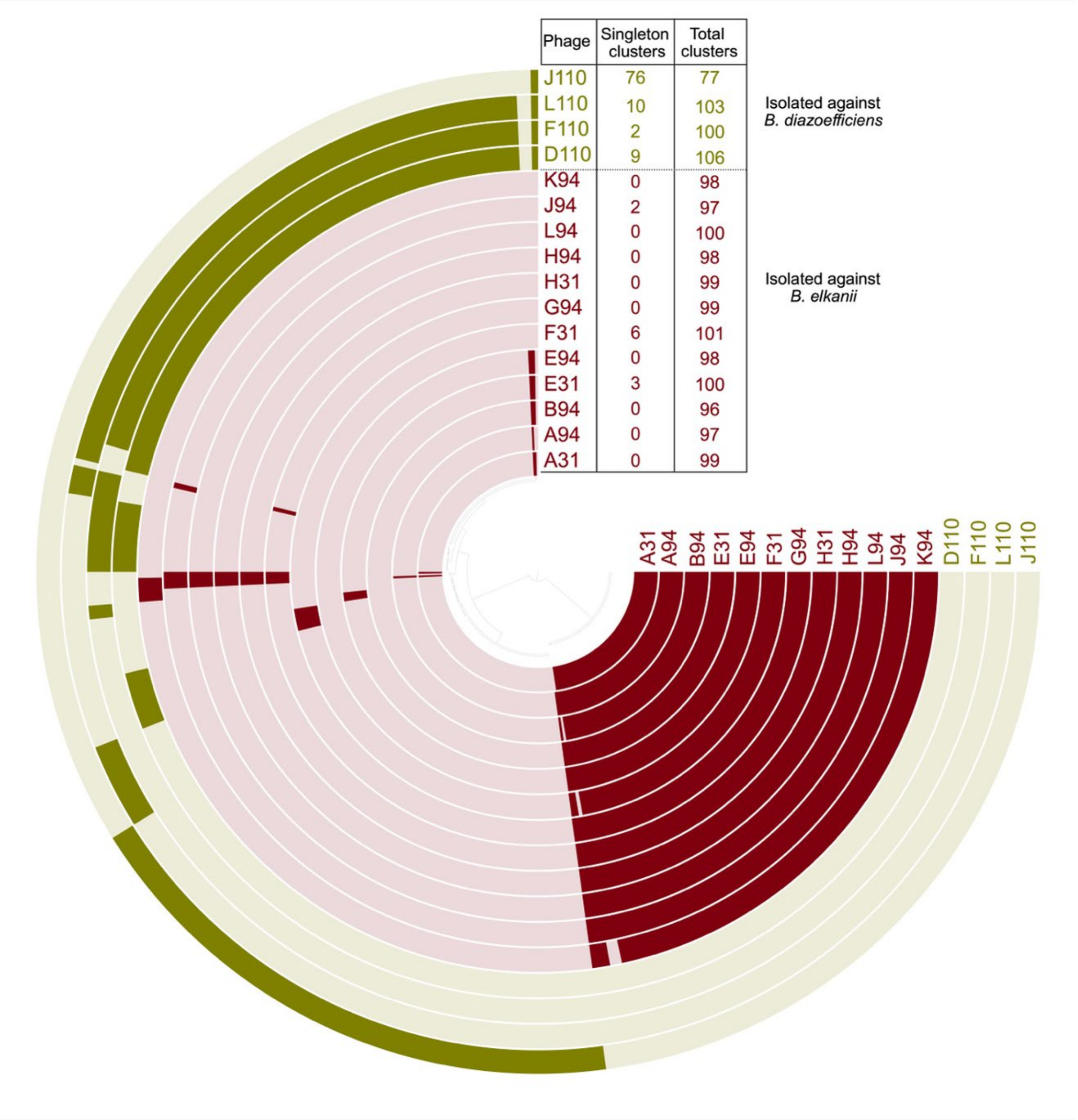


**Figure S4.** Pangenome display of 16 phages virulent on soybean *Bradyrhizobium* spp. isolated from Delaware soils. The pangenome clusters based on presence/absence across the 16 genomes (Euclidean distance; Ward linkage). The center tree arranges the gene clusters identified across the genomes. Each concentric layer represents a phage genome, with dark shading indicating the presence of a gene cluster in that genome, and light shading representing the absence of a gene cluster. The positioning of the gene clusters does not represent their order in the genomes. Layers are grouped and color-coded by the original isolation host species: *B. elkanii* (red), *B. diazoefficiens* (green). This figure highlights the core, accessory, and unique gene clusters, showing both the conservation and variability across the three phage populations.
