## Supplemental table 1 for "Comparative Analysis Reveals Host Species-Dependent Diversity Among 16 Virulent Bacteriophages Isolated Against Soybean *Bradyrhizobium* spp"

**Table S1.** Means and corresponding standard deviations of morphological characteristics for 16 virulent *Bradyrhizobium* phages isolated from Delaware soils. Individual phages were measured for capsid diameter, capsid length, tail diameter, and tail length. Capsid volumes were calculated using a formula for the volume of an ellipsoid.

| Isolation species  and strain | Sample size | Capsid  diameter  (nm) | Capsid  length  (nm) | Capsid  volume  (nm^3^) | Tail  diameter (nm) | Tail   length (nm) |
| --- | --- | --- | --- | --- | --- | --- |
| *B. elkanii*  USDA 31 | 62 | 58.1 ± 5.11 | 71.5 ± 7.27 | 1.29E^5^ | 12.9 ± 0.48 | 128.3 ± 3.18 |
| *B. elkanii*  USDA 94 | 134 | 59.3 ± 3.50 | 72.2 ± 4.84 | 1.33E^5^ | 12.3 ± 1.04 | 126.8 ± 4.87 |
| Mean values for  *B. elkanii* | 196 | 58.9±4.07 | 71.9±5.66 | 1.32E^5^ | 12.5 ± 2.05 | 127.1 ± 11.96 |
| *B. diazoefficiens*  USDA 110 | 113 | 67.3 ± 5.85 | 67.9 ± 4.69 | 1.63E^5^ | nd^1^ | 16.6 ± 2.45^2^ |

^1^nd, not determined. Phages isolated against USDA 110 displayed a podophage-like morphology that prevented reliable tail diameter measurements.

^2^Tail length is not representative of the full sample size due to unreliable tail measurements for some images.
