## Supplemental table 2 for "Comparative Analysis Reveals Host Species-Dependent Diversity Among 16 Virulent Bacteriophages Isolated Against Soybean *Bradyrhizobium* spp"

**Table S2**. Genomic features of 16 virulent *Bradyrhizobium* phages isolated from Delaware soils. Genome length in base pairs, percent GC content, total number of genes, and the number of functionally annotated genes are shown.

| Isolation host  species | Lytic phage  isolate | Total genome  length (bp) | GC content  (%) | Total number  of genes^1^ | Genes with functional  annotations^1^ (%) |
| --- | --- | --- | --- | --- | --- |
|  | A31 | 63,461 | 67.0 | 101 | 24.75 |
|  | E31 | 62,868 | 67.1 | 99 | 25.25 |
|  | F31 | 63,113 | 67.1 | 101 | 24.75 |
|  | H31 | 63,205 | 66.9 | 102 | 24.51 |
|  | A94 | 62,724 | 67.0 | 99 | 25.25 |
| *B. elkanii* | B94 | 62,580 | 67.1 | 99 | 25.25 |
|  | E94 | 62,507 | 67.1 | 99 | 25.25 |
|  | G94 | 63,251 | 66.9 | 100 | 25.00 |
|  | H94 | 63,205 | 67.0 | 100 | 25.00 |
|  | J94 | 64,087 | 66.7 | 103 | 24.27 |
|  | K94 | 62,466 | 66.9 | 99 | 24.24 |
|  | L94 | 63,206 | 67.0 | 97 | 25.77 |
|  | D110^2^ | 57,535 | 43.0 | 105 | 21.90 |
| *B. diazoefficiens* | F110^3^ | 55,636 | 43.5 | 98 | 20.41 |
|  | L110^2^ | 57,005 | 44.2 | 105 | 21.90 |
|  | J110 | 41,973 | 58.9 | 78 | 19.23 |

^1^Includes tRNA and tmRNA genes

^2^Genome contains tRNAs: tRNA-Leu (taa), tRNA-Thr (tgt)

^3^Genome contains tRNAs: tRNA-Leu (taa), tRNA-Thr (tgt), and tRNA-Met (cat)
